# Asymmetric Engagement of Dimeric CRL3^KBTBD4^ by the Molecular Glue UM171 Licenses Degradation of HDAC1/2 Complexes

**DOI:** 10.1101/2024.05.14.593897

**Authors:** Megan J.R. Yeo, Olivia Zhang, Xiaowen Xie, Eunju Nam, N. Connor Payne, Pallavi M. Gosavi, Hui Si Kwok, Irtiza Iram, Ceejay Lee, Jiaming Li, Nicholas J. Chen, Hanjie Jiang, Zhipeng A. Wang, Kwangwoon Lee, Haibin Mao, Stefan A. Harry, Idris A. Barakat, Mariko Takahashi, Amanda L. Waterbury, Marco Barone, Andrea Mattevi, Liron Bar-Peled, Philip A. Cole, Ralph Mazitschek, Brian B. Liau, Ning Zheng

**Affiliations:** Department of Chemistry and Chemical Biology, Harvard University, Cambridge, MA, USA; Broad Institute of MIT and Harvard, Cambridge, MA, USA; Department of Pharmacology University of Washington, Seattle, WA, USA; Howard Hughes Medical Institute, University of Washington, Seattle, WA, USA; Division of Genetics, Department of Medicine, Brigham and Women’s Hospital; Department of Biological Chemistry and Molecular Pharmacology, Harvard Medical School, Boston, MA, USA; Center for Systems Biology, Massachusetts General Hospital, Boston, MA, USA; Harvard T.H. Chan School of Public Health, Boston, MA, USA; Krantz Family Center for Cancer Research, Massachusetts General Hospital, Boston MA, USA; Department of Biology and Biotechnology Lazzaro Spallanzani, University of Pavia, Pavia, Italy; Department of Medicine, Harvard Medical School, Boston, MA, USA

## Abstract

UM171 is a potent small molecule agonist of *ex vivo* human hematopoietic stem cell (HSC) self-renewal^1^, a process that is tightly controlled by epigenetic regulation. By co-opting KBTBD4, a substrate receptor of the CULLIN3-RING E3 ubiquitin ligase complex, UM171 promotes the degradation of members of the CoREST transcriptional corepressor complex, thereby limiting HSC attrition^2,3^. However, the direct target and mechanism of action of UM171 remain unclear. Here, we reveal that UM171 acts as a molecular glue to induce high-affinity interactions between KBTBD4 and HDAC1 to promote the degradation of select HDAC1/2 corepressor complexes. Through proteomics and chemical inhibitor studies, we discover that the principal target of UM171 is HDAC1/2. Cryo-electron microscopy (cryo-EM) analysis of dimeric KBTBD4 bound to UM171 and the LSD1-HDAC1-CoREST complex unveils an unexpected asymmetric assembly, in which a single UM171 molecule enables a pair of KBTBD4 KELCH-repeat propeller domains to recruit HDAC1 by clamping on its catalytic domain. One of the KBTBD4 propellers partially masks the rim of the HDAC1 active site pocket, which is exploited by UM171 to extend the E3-neo-substrate interface. The other propeller cooperatively strengthens HDAC1 binding via a separate and distinct interface. The overall neomorphic interaction is further buttressed by an endogenous cofactor of HDAC1-CoREST, inositol hexakisphosphate, which makes direct contacts with KBTBD4 and acts as a second molecular glue. The functional relevance of the quaternary complex interaction surfaces defined by cryo-EM is demonstrated by in situ base editor scanning of KBTBD4 and HDAC1. By delineating the direct target of UM171 and its mechanism of action, our results reveal how the cooperativity offered by a large dimeric CRL E3 family can be leveraged by a small molecule degrader and establish for the first time a dual molecular glue paradigm.

## Introduction

Molecular glue degraders are small molecules capable of promoting the ubiquitination and degradation of proteins, including those conventionally deemed undruggable^4,5^. Unlike bifunctional proteolysis targeting chimeras (PROTACs), molecular glue degraders induce high-affinity interactions between an E3 ubiquitin ligase and a substrate protein without showing a detectable affinity to at least one of the protein partners^6^. While rapid progress has been made in the rational design and clinical trials of PROTACs, the development of molecular glue degraders has been protracted due to poor understanding of their functional prerequisites and capacities. The plant hormones auxin and jasmonate are the first documented molecular glue degraders, which target the F-box proteins, the substrate receptors of CULLIN1-RING ligase (CRL1) complexes^7–9^. In human cells, the best characterized molecular glue degraders include thalidomide and its derivatives, aryl-sulfonamides, and CDK12 inhibitors^4^. Curiously, these synthetic compounds all co-opt the CULLIN4-RING ligases (CRL4s), raising the question of whether other ubiquitin ligases can be reprogrammed by molecular glue degraders. CRL3s, in particular, represent the largest family of CRLs with nearly 200 substrate receptors. Its family members, such as CRL3^KEAP1^ and CRL3^SPOP^, as well as select members of other CRL complexes, including CRL1^FBXW7^ and CRL1^β-TrCP^, form constitutive homodimers that are exploited by endogenous substrates for cooperative binding^10–14^. Whether these E3 ligases can be leveraged by molecular glue degraders, especially in ways to exploit their intrinsic cooperativity, remains uncertain. Despite the promise of targeted protein degradation as a modality for drug discovery, the scarcity of molecular glue degraders with high potency and efficacy has restricted our insights into their activity requirements, target and E3 scaffold preferences, and functional versatility.

Small molecule agonists that promote the self-renewal and expansion of HSCs are important reagents for basic research and have clinical applications for cell-based therapies^15,16^. UM171 (**Fig. 1a**) was originally identified in a phenotypic screen for molecules promoting HSC expansion^1^. Despite its wide use and progression into human clinical trials, the precise mechanism and direct target of UM171 have eluded the field for a decade. Recently, UM171 was shown to induce widespread activation of enhancers and transcriptional programs through degradation of key subunits of the CoREST corepressor complex^2,3^. The CoREST core complex comprises lysine-specific histone demethylase 1a (LSD1), CoREST, and either the histone deacetylase HDAC1 or its paralog HDAC2. In the LSD1-HDAC1/2-CoREST (LHC) complex, CoREST serves as a scaffold to recruit LSD1 and HDAC1/2 at its two ends^17^. Upon addition of UM171, CoREST is ubiquitinated and rapidly degraded, followed shortly by subsequent degradation of LSD1^2^, whose stability is dependent on CoREST^18^. CoREST degradation is mediated by KBTBD4^2^, a BTB-KELCH E3 substrate adaptor belonging to the CULLIN3-RING ligase (CRL3) family. Here, we identify HDAC1/2 as the crucial target of UM171 for KBTBD4-mediated CoREST degradation and elucidate the mechanism of action of UM171 in synergy with the HDAC1/2 cofactor inositol hexakisphosphate^19,20^. Our findings demonstrate the capacity of a molecular glue degrader duo to exploit a member within the largest family of CRL E3 ligases by co-opting an asymmetric assembly of the homodimeric E3 scaffold.

**Figure 1.**
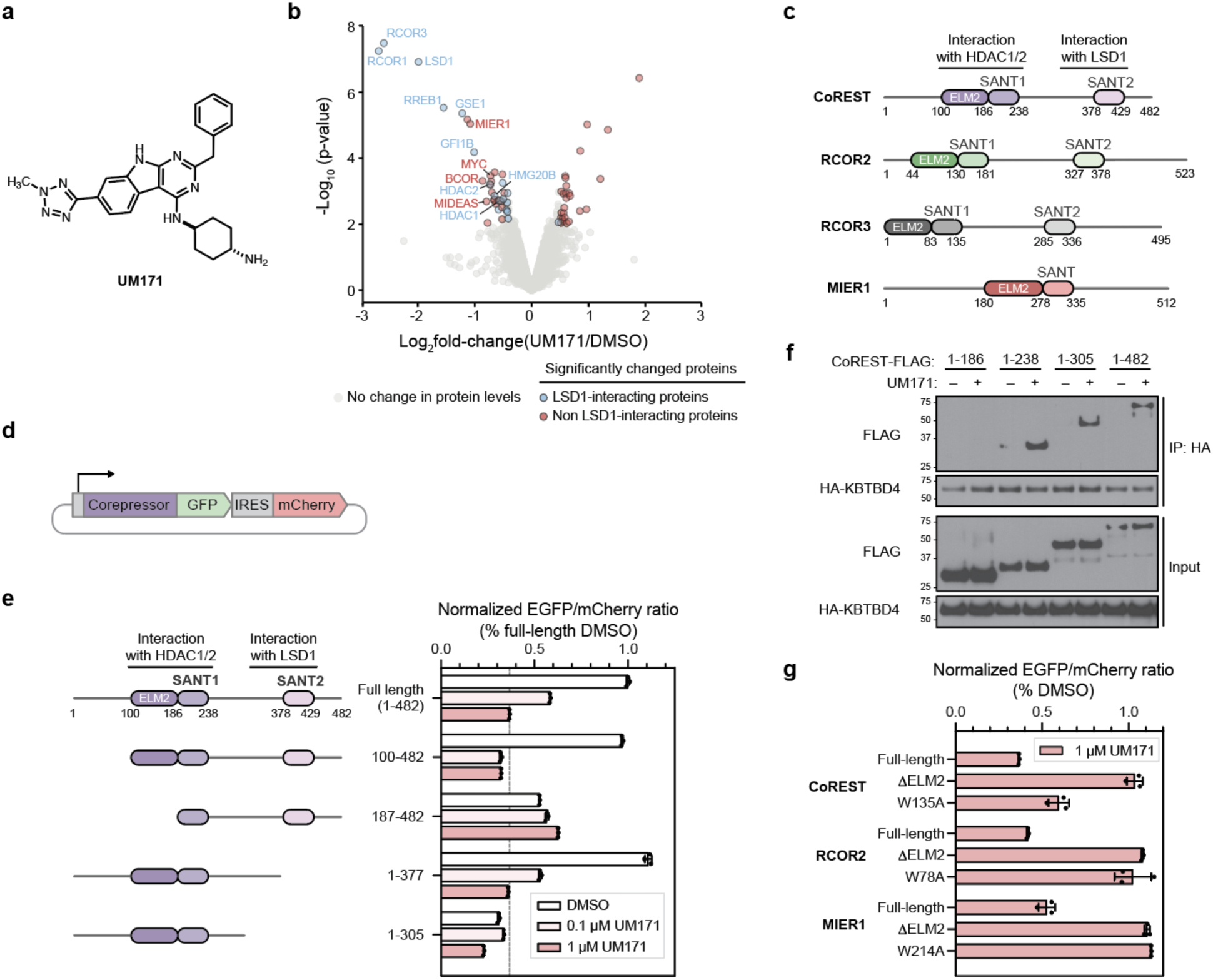
UM171-induced degradation of CoREST depends on HDAC1/2 interaction. **a)** Chemical structure of UM171. **b)** Volcano plot showing proteins up- and down-regulated upon UM171 treatment in SET-2 cells. Colored dots (blue, red) show proteins with | log_2_(fold-change) | > 0.5 in UM171 versus DMSO treatment and p-value < 0.01. Dots in red and blue depict proteins enriched or absent in LSD1 co-IP/MS, respectively. **c)** Protein domain maps of CoREST, RCOR2, RCOR3, and MIER1. **d)** Schematic of corepressor-GFP/mCherry stability reporter. **e)** GFP to mCherry fluorescence ratio measured by flow cytometry of MOLM-13 cells expressing the indicated CoREST-GFP reporter treated with DMSO or UM171 (0.1 or 1 µM) for 24 h. Bars represent mean ± s.d. of *n = 3* replicates. **f)** Immunoblots of HA IP in the presence of 1 µM UM171 (1 h) or DMSO and 1 µM MLN4924 (3 h) from 293T cells transfected with HA-KBTBD4 and the indicated CoREST-FLAG construct. **g)** GFP to mCherry fluorescence ratio measured by flow cytometry of MOLM-13 cells expressing the indicated corepressor-GFP reporter treated with UM171 (1 µM) for 24 h. Bars represent mean ± s.d. of *n = 3* replicates. Results in **e-g** are representative of two independent experiments.

### UM171 promotes selective degradation of HDAC1/2 complexes

The direct binding target of UM171 has been elusive for a decade. To resolve the mystery, we first sought to define the protein substrates depleted by UM171 treatment. To determine the full repertoire of proteins degraded by UM171, we conducted a global proteomics analysis in two UM171-sensitive leukemia cell lines, SET-2 and MV4;11, after vehicle or UM171 treatment (6 h, 1 µM). LSD1 and two CoREST homologs, RCOR1 (hereafter referred to as CoREST) and RCOR3, were the most significantly depleted proteins after UM171 treatment (**Fig. 1b; Extended Data Fig. 1a**). RCOR2, another CoREST homolog, is not expressed in these AML cell lines^21^. All three CoREST homologs share conserved ELM2 and SANT1/2 domains as well as a linker region that forms a highly stable complex with LSD1 (**Fig. 1c**)^18,22^. Several other highly down-regulated proteins are components of the broader LHC complex (e.g., RREB1, GSE1, HMG20B, GFI1B), suggesting extensive collateral degradation as previously observed with HDAC-targeting PROTACs^23^. To assess which down-regulated proteins are direct versus collateral targets of KBTBD4, we identified LSD1-interacting proteins using co-immunoprecipitation (co-IP) MS in SET-2 cells (**Extended Data Fig. 1b**). As expected, many proteins depleted by UM171 treatment associate with LHC (blue dots, **Fig. 1b; Extended Data Fig. 1a, b**). However, MIER1 is highly depleted but does not co-IP with LSD1 (red dots), indicating that it might be an alternative substrate of UM171-KBTBD4 and that LSD1 is not the direct target of UM171. Notably, all three CoREST homologs, MIER1, as well as MIDEAS — which is depleted by UM171 to a lesser extent — contain an ELM2-SANT tandem domain (**Fig. 1c**)^20^. Collectively, these results demonstrate that UM171 promotes selective degradation of several corepressors characterized by an ELM2-SANT domain.

We next defined the region(s) of the corepressors necessary for UM171-induced degradation by using a fluorescent degrader reporter system^24^, in which full-length or truncated corepressor variants are fused in-frame with GFP followed by an internal ribosome entry site (IRES) and mCherry (**Fig. 1d; Extended Data Fig. 2a**). We started with CoREST since it is most potently degraded. While deletion of the N-terminal 1-100 amino acids (aa) or the C-terminal SANT2 domain (aa 377-482) had minimal impact, removal of aa 1-186 completely blocked CoREST-GFP degradation by UM171 (**Fig. 1e; Extended Data Fig. 1c**), providing evidence that the ELM2 domain is necessary. A larger C-terminal deletion reduced baseline stability of CoREST-GFP, likely due to an inability to complex with LSD1^18,22^. Nonetheless, UM171 treatment still decreased CoREST(aa 1-305)-GFP levels, showing that LSD1 is not necessary for CoREST degradation. Co-IP experiments demonstrated that CoREST(aa 1-238)-FLAG is sufficient to interact with HA-KBTBD4 while CoREST(aa 1-186)-FLAG cannot (**Fig. 1f**). Lastly, using the GFP-reporter system, we observed that the ELM2-SANT domains were also required for UM171-induced degradation of MIER1 and RCOR2 (**Fig. 1g**). Altogether, these results show that the ELM2-SANT domains are necessary for UM171-induced degradation of corepressors.

In all the tested corepressors, the ELM2-SANT tandem domain mediates complexation with HDAC1 and its paralog HDAC2 interchangeably^20^ (**Fig. 1c**). Hence, we reasoned that not only the ELM2-SANT domain but also HDAC1/2 might be necessary for corepressor degradation by UM171. Mutation of MIER1 Trp214 to alanine (W214A) has been previously shown to abrogate HDAC1/2 binding to MIER1 by disrupting the ELM2-HDAC1 interface^25^. We found that MIER1 W214A and the analogous Trp to Ala mutants of CoREST (W135A) and RCOR2 (W78A) exhibited significantly decreased UM171-induced degradation (**Fig. 1g**), supporting the notion that interaction with HDAC1/2 is necessary. Altogether, our findings demonstrate that the ELM2-SANT tandem domain and HDAC1/2 are critical for UM171/KBTBD4-mediated degradation of select corepressor complexes.

### HDAC1/2 mediates LHC-KBTBD4 ternary complex formation

We further investigated the mechanistic involvement of HDAC1/2 in UM171’s mechanism of action by assessing endogenous CoREST degradation in *HDAC1* or *HDAC2* knockout cell lines. To do so, we first engineered a K562 knock-in cell line with GFP fused to the C-terminus of endogenous CoREST. Treatment of these cells with UM171 led to rapid KBTBD4-dependent CoREST-GFP depletion (**Extended Data Fig. 3a, b**). In this system, knockout (KO) of *HDAC1* or *HDAC2* partially blocked CoREST-GFP depletion by UM171, likely due to functional redundancy between the two paralogs^26^ (**Fig. 2a; Extended Data Fig. 3c**). Notably, pre-treatment of cells with HDAC active-site inhibitors^27^ — including suberoylanilide hydroxamic acid (SAHA, also known as vorinostat), CI-994, and Cpd-60^28^ — also blocked CoREST-GFP degradation as well as co-IP of FLAG-KBTBD4 with CoREST induced by UM171 (**Fig. 2b, c**). However, UM171 had no impact on recombinant HDAC1 and HDAC2 enzymatic activity in biochemical assays (**Extended Data Fig. 3d**). Altogether, these data demonstrate that HDAC1/2 and their accessible active sites are required for UM171-induced degradation of CoREST.

**Figure 2.**
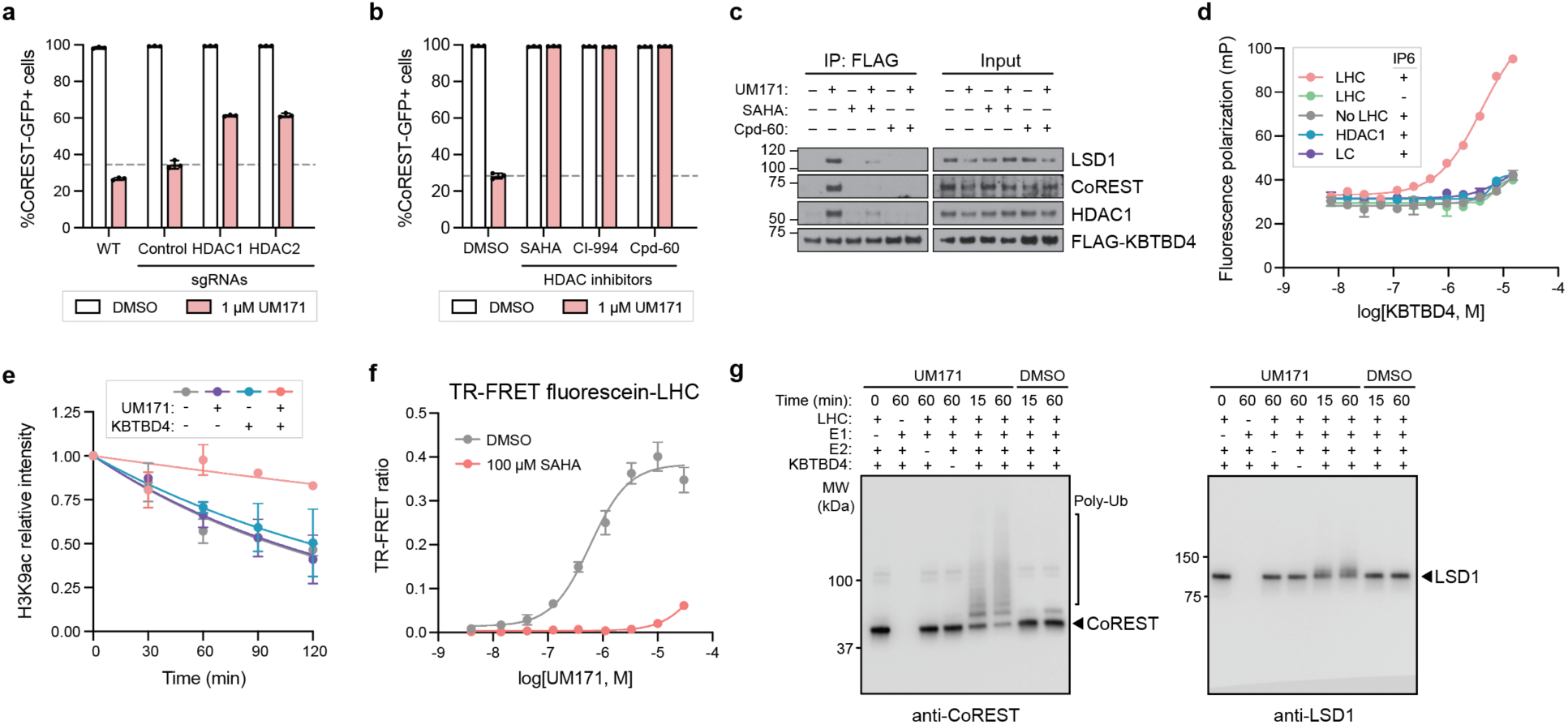
HDAC1 mediates LHC-UM171-KBTBD4 ternary complex formation. **a)** Flow cytometry quantification of K562 CoREST-GFP cells transduced with indicated sgRNAs for treatment with DMSO or UM171 (1 µM) for 24 h. Bars represent mean ± s.d. of *n = 3* replicates. **b)** Flow cytometry quantification of K562 CoREST-GFP cells for co-treatment with indicated HDAC inhibitors (10 µM) and UM171 (1 µM) for 24 h. Bars represent mean ± s.d. of *n = 3* replicates. **c)** Immunoblots of FLAG IP in the presence of 5 µM UM171, 10 µM HDACi, or vehicle, and 1 µM MLN4924 from K562 cells stably expressing FLAG-KBTBD4. **d)** Dose-response curve showing FP of JL1 (10 nM) (*y*-axis) with KBTBD4 (*x*-axis) in the presence or absence of LHC (20 nM). Data are mean ± s.d. of *n* = 3 replicates. **e)** Line plots showing quantification of LHC (90 nM) deacetylase activity on H3K9ac-modified mononucleosomes in the presence or absence of UM171 (10 µM) and/or KBTBD4 (300 nM). Data are mean ± s.e.m. of *n =* 2 replicates. **f)** Dose-response curve showing TR-FRET signal (*y*-axis) between fluorescein-LHC (40 nM) and anti-His CoraFluor-1-labeled antibody with His-KBTBD4 (40 nM) in the presence of varying concentrations of UM171 (*x*-axis). Data are mean ± s.d. of *n* = 2 replicates. **g)** Immunoblots for CoREST (left) and LSD1 (right) of in vitro ubiquitination assays of CRL3^KBTBD4^ (500 nM) with fluorescein-LHC (500 nM) in the presence of DMSO or UM171 (10 µM). Results in **a-f** are representative of two independent experiments. Results of **g** are representative of three independent experiments.

We next sought to determine if UM171 is sufficient to induce ternary complex formation with KBTBD4 and members of the LHC complex. His-KBTBD4 was expressed and purified from Sf9 cells and LHC containing HDAC1 was prepared as previously described (see Methods)^17,29^. Fluorescence polarization (FP) assays using a derivative of UM171 conjugated to tetramethylrhodamine, **JL1**, showed binding of **JL1** only in the presence of both KBTBD4 and LHC together, and furthermore only in the presence of inositol hexakisphosphate (InsP_6_) (**Fig. 2d; Extended Data Fig. 3e, f**). InsP_6_ has been previously shown to stabilize the interaction of HDAC1/2 with their cognate corepressors^19,20,30^. Accordingly, all subsequent experiments were conducted with 50 µM InsP_6_ unless otherwise noted. **JL1**-KBTBD4 binding was not observed with LSD1-CoREST or HDAC1 alone, showing that the full LHC complex is necessary for ternary complex formation (**Fig. 2d**), consistent with microscale thermophoresis (MST) assays (**Extended Data Fig. 3g, h**). Moreover, addition of SAHA dose-dependently blocked FP with an IC_50_ comparable to its affinity for HDAC1/2 and LHC^30^, which is consistent with our degradation experiments (**Fig. 2b, Extended Data Fig. 3i**). Collectively, these results demonstrate that UM171 exhibits highly cooperative binding and does not exhibit strong affinity to either KBTBD4 or LHC alone. Consistent with this notion, UM171 could only inhibit LHC deacetylase activity on recombinant nucleosomes in the presence of KBTBD4 (**Fig. 2e; Extended Data Fig. 3j**), further suggesting that complexation with the E3 obstructs the HDAC1 active site.

To directly assess the association between KBTBD4 and LHC, we next used time-resolved Förster resonance energy transfer (TR-FRET) with labeled protein complexes. To conduct these assays, an ectopic cysteine residue was introduced at the N-terminus of CoREST (aa 84-482) and selectively labeled with fluorescein (see Methods) while His-KBTBD4 was labeled in situ with an anti-His CoraFluor-1 antibody^30,31^. UM171 induced strong TR-FRET signal in a dose-dependent manner, indicating association between fluorescein-LHC and His-KBTBD4 with an apparent EC_50_ value of 542 nM under the experimental conditions (**Fig. 2f**). As expected, co-treatment with SAHA blocked UM171-induced LHC-KBTBD4 association. Dose-response titration of fluorescein-LHC against His-KBTBD4 in the presence of UM171 and InsP_6_ at saturating concentrations yielded an equilibrium dissociation constant (*K*_D_) value of 16 nM for the UM171-mediated LHC-KBTBD4 interactions (**Extended Data Fig. 3k**). Lastly, we established that reconstituted CRL3^KBTBD4^ is sufficient to mediate ubiquitination of LHC in vitro, which was significantly potentiated by addition of UM171 (**Fig. 2g; Extended Data Fig. 3l**). Collectively, these results demonstrate that UM171 exhibits highly cooperative binding and possibly acts as a molecular glue to stabilize a ternary complex with KBTBD4 and LHC. Importantly, we establish the critical roles of HDAC1/2 and InsP_6_ in mediating UM171-induced ternary complex formation, defining the minimal components necessary to reconstitute the complex for structural analysis.

### Overall structure of the KBTBD4-UM171-LHC complex

To resolve the mechanism of action of UM171, we next assembled and isolated the KBTBD4-LHC-UM171 complex by size exclusion chromatography in the presence of InsP_6_ to maintain the integrity of the complex. For this purpose, we employed an N-terminally 10 aa deleted version of LSD1, which enhances the production and stability of LHC. Single particle cryo-EM analysis of the KBTBD4-LHC-UM171 complex yielded a highly interpretable map at an average resolution of 3.93 Å (**Extended Data Fig. 4, Extended Data Table 1**). In the 3D reconstruction map of the KBTBD4-LHC complex, KBTBD4 and HDAC1 are well resolved, whereas only partial densities are visible for the ELM2-SANT1 domain of CoREST. The rest of LHC could not be resolved, most likely due to the flexible nature of CoREST.

The KBTBD4-LHC complex adopts an unexpected asymmetric architecture, in which two protomers of a KBTBD4 homodimer, hereafter referred to as KBTBD4-A and KBTBD4-B, simultaneously engage a single copy of HDAC1-CoREST in a bidentate fashion (**Fig. 3a**). Although the SANT1 domain of CoREST is brought into close vicinity of KBTBD4, the closest Cα atoms of the two proteins remain 9 Å away from each other, and therefore the interactions between KBTBD4 and HDAC1 exclusively comprise the contacts made by the E3 to LHC. In the asymmetric assembly, the two KBTBD4 protomers interact with HDAC1 via two separate interfaces: (1) KBTBD4-A engages with the outer edge of the HDAC1 catalytic domain, whereas (2) KBTBD4-B cups HDAC1 by packing against the rim of its active site pocket.

**Figure 3.**
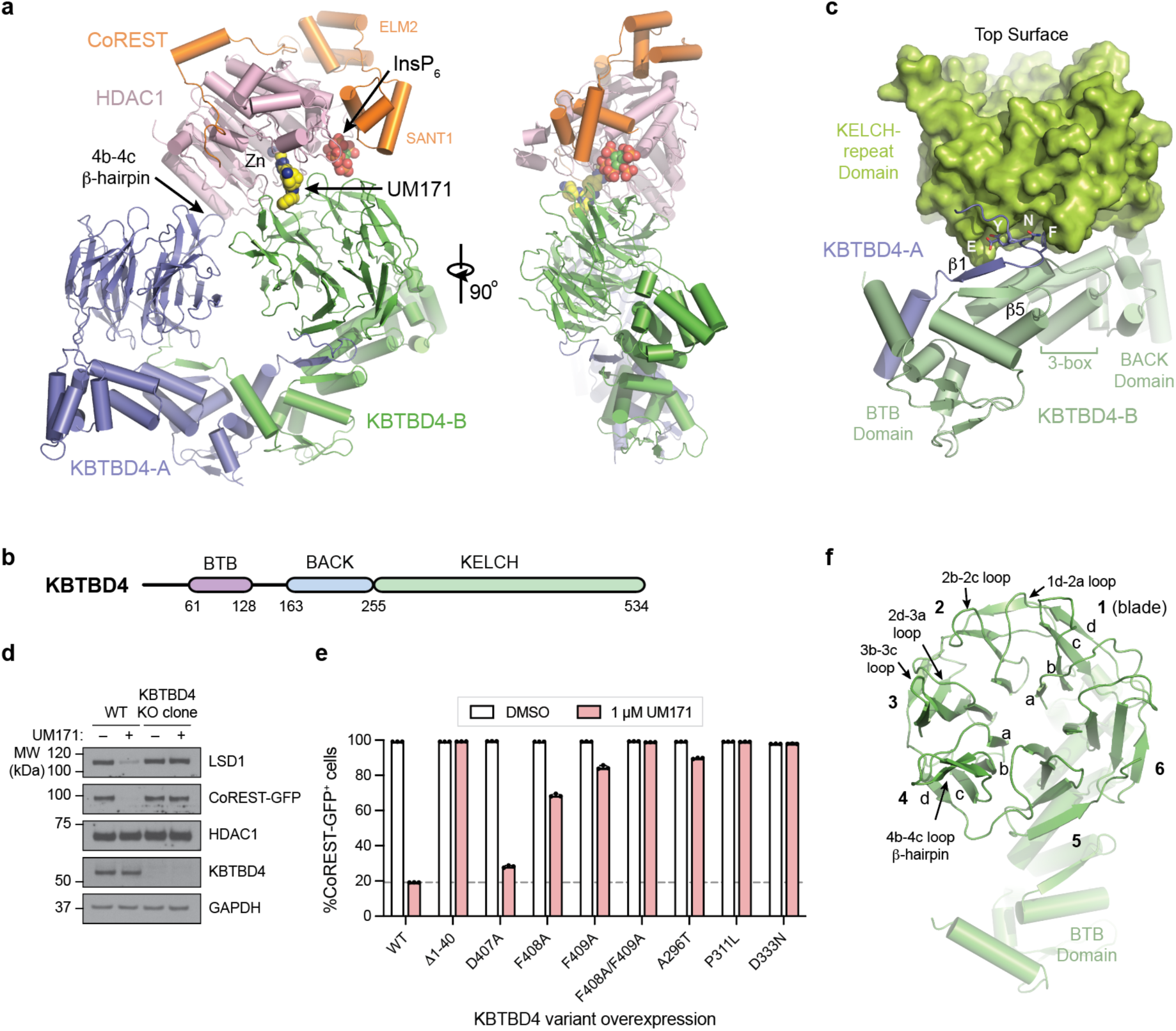
Overall structure of KBTBD4-HDAC1-CoREST-UM171 complex. **a)** Two orthogonal views of HDAC1 (pink) and CoREST (orange)-bound KBTBD4 (green/blue) with UM171 (space-filling model in yellow and blue) and inositol hexakisphosphate, InsP_6_ (space-filling model in green and red). **b)** Schematic showing protein domains of KBTBD4. **c)** The BTB and BACK domains in cartoon representation and the KELCH-repeat domain in surface representation of a KBTBD4 protomer (green) in the KBTBD4 dimer. The N-terminal region of the other protomer with the domain-swapped β1-strand flanked by the EYNF motif and an α-helix is shown in blue. Helices predicted to bind CUL3 is indicated as 3-box. **d)** Immunoblot showing LSD1, CoREST-GFP, HDAC1, KBTBD4, and GAPDH in wild-type K562 CoREST-GFP knock-in cells and a clonal cell line with KBTBD4 knockout in the presence or absence of UM171 (1 µM, 6 h). **e)** Flow cytometry quantification of K562 KBTBD4-null CoREST-GFP cells overexpressing indicated KBTBD4 variants after treatment with DMSO or UM171 (1 µM) for 24 h. Bars represent mean ± s.d. of *n = 3* replicates. **f)** The KELCH-repeat β-propeller domain of KBTBD4 with its secondary structure elements annotated. Results in **d** and **e** are representative of two independent experiments.

As the hallmark of the E3-neo-substrate complex, a single molecule of UM171 is situated at the HDAC1-KBTBD4-B interface. The *N-*methyl-tetrazole of the compound is inserted into the outer periphery of the active site pocket of the deacetylase, while the tricyclic pyrimidoindole core is embedded in an induced surface groove of KBTBD4-B (**Fig. 3a**). By interacting with both HDAC1 and KBTBD4-B, UM171 fills an exposed gap between the two proteins with exquisite shape complementarity, extending the protein-protein interaction interface as a molecular glue. Directly adjacent to the UM171 binding site, InsP_6_ is nestled at the three-protein junction, making direct contacts with HDAC1, CoREST, and KBTBD4-B to stabilize the E3-neo-substrate complex as a second molecular glue. Together, the E3 ligase dimer, the neo-substrate complex, and the two small molecules bury a total surface area of ∼ 2,300 Å^2^, with more than half of the interfaces contributed by protein-protein interactions.

### Asymmetric arrangement of KBTBD4 protomers

KBTBD4 contains an N-terminal BTB domain, a central BACK domain, and a C-terminal KELCH-repeat propeller domain (**Fig. 3b**). As expected, KBTBD4 forms a homodimer via its BTB domain, which is characterized by a domain-swapped two-stranded β-sheet (β1 & β5) as part of the dimer interface (**Fig. 3c**)^32^. The predicted CUL3-binding 3-box helices of the KBTBD4 BTB domain are extended by five short helices in the BACK domain, which is connected to the KELCH-repeat domain via a linker sequence^12,33^. Unique to KBTBD4 among other BTB and KELCH domain-containing proteins, the N-terminal β1 strand of its BTB domain is led by a highly conserved ENYF motif (**Extended Data Fig. 5**), which packs against the C-terminal KELCH-repeat propeller of the second protomer upon domain swapping. This motif appears to structurally couple the two halves of the CRL3 substrate receptor (**Fig. 3c**). Overexpression of a KBTBD4 mutant lacking the ENYF motif (KBTBD4 Δ1-40) in CoREST-GFP KBTBD4-null cells completely abrogates CoREST degradation by UM171 (**Fig. 3d, e**), suggesting that proper positioning of the two KELCH-repeat domains against the BTB domains in the dimer is critical for E3 function.

The KBTBD4 dimer has an overall twofold symmetry with its two protomers superimposable with an R.M.S.D. of 0.85 Å over 445 Cα atoms (**Extended Data Fig. 6a, b**). Outside the N-terminal BTB domain, the two KBTBD4 protomers do not physically contact one another. The C-terminal KELCH-repeat domain adopts a canonical six-bladed propeller fold (**Fig. 3f**). From the center to the outer edge of the propeller, the four β-strands within a blade are conventionally named “a” to “d”^34^. Distinct from the other five blades, the fourth blade of the KBTBD4 KELCH-repeat propeller features an extra-long b-c loop, which adopts a β-hairpin structure that protrudes from the top surface of the propeller (**Fig. 3f**). Although the central pocket presented by the top surface of a propeller fold is frequently used by KELCH-repeat domain-containing E3s to engage their substrates^35–37^, the two propellers in the KBTBD4 dimer orient their top surfaces to opposite directions (**Extended Data Fig. 6a**) and instead use mostly their lateral surfaces and the b-c loops to simultaneously recognize HDAC1.

### Interface between HDAC1 and KBTBD4-A

HDAC1 adopts a single α/β fold with a central eight-stranded parallel β-sheet sandwiched by α-helices on its two faces^38^. The active site is characterized by a catalytic zinc ion deep in the narrow pocket and an outer rim involved in recognition of its histone substrates. Superposition of the MTA1-bound and KBTBD4-bound HDAC1 structures shows that the deacetylase does not undergo major conformational changes upon binding to the E3 dimer in the presence of UM171 (**Extended Data Fig. 6c**)^39^. Nevertheless, an α-helix and a loop region concealing the C-terminal edge of HDAC1’s central β-sheet are slightly spread apart by KBTBD4-A to promote KBTBD4-LHC-UM171 complex formation (**Extended Data Fig. 6d**).

At this interface, the β-hairpin of the KBTBD4-A 4b-4c loop wedges into a hydrophobic cleft demarcated by the outer strand (β6) of HDAC1’s central β-sheet and its two surrounding secondary structure elements. Two adjacent phenylalanine residues (Phe408 and Phe409) at the tip of the KBTBD4-A β-hairpin make close contacts with five hydrophobic residues in HDAC1 (Tyr201, Leu211, Pro227, Tyr358, and Ile362) (**Fig. 4a**), which are conserved in HDAC2. These interactions are reinforced by an intermolecular salt bridge formed between Asp407 of KBTBD4-A and Arg229 of HDAC1. Three additional positively charged residues of the deacetylase (Arg212, Lys361, and Arg365) further substantiate the interface by packing their aliphatic side chains against the two KBTBD4-A phenylalanine residues. Consistent with an important role in stabilizing the KBTBD4-LHC-UM171 complex, mutation of either of the two KBTBD4 phenylalanine residues (Phe408, Phe409) to alanine blocked CoREST degradation by UM171 (**Fig. 3e**). By contrast, D407A had a lesser impact on CoREST degradation. Altogether, these data support the notion that engagement by KBTBD4-A is critical for HDAC1 recognition and subsequent CoREST degradation.

**Figure 4.**
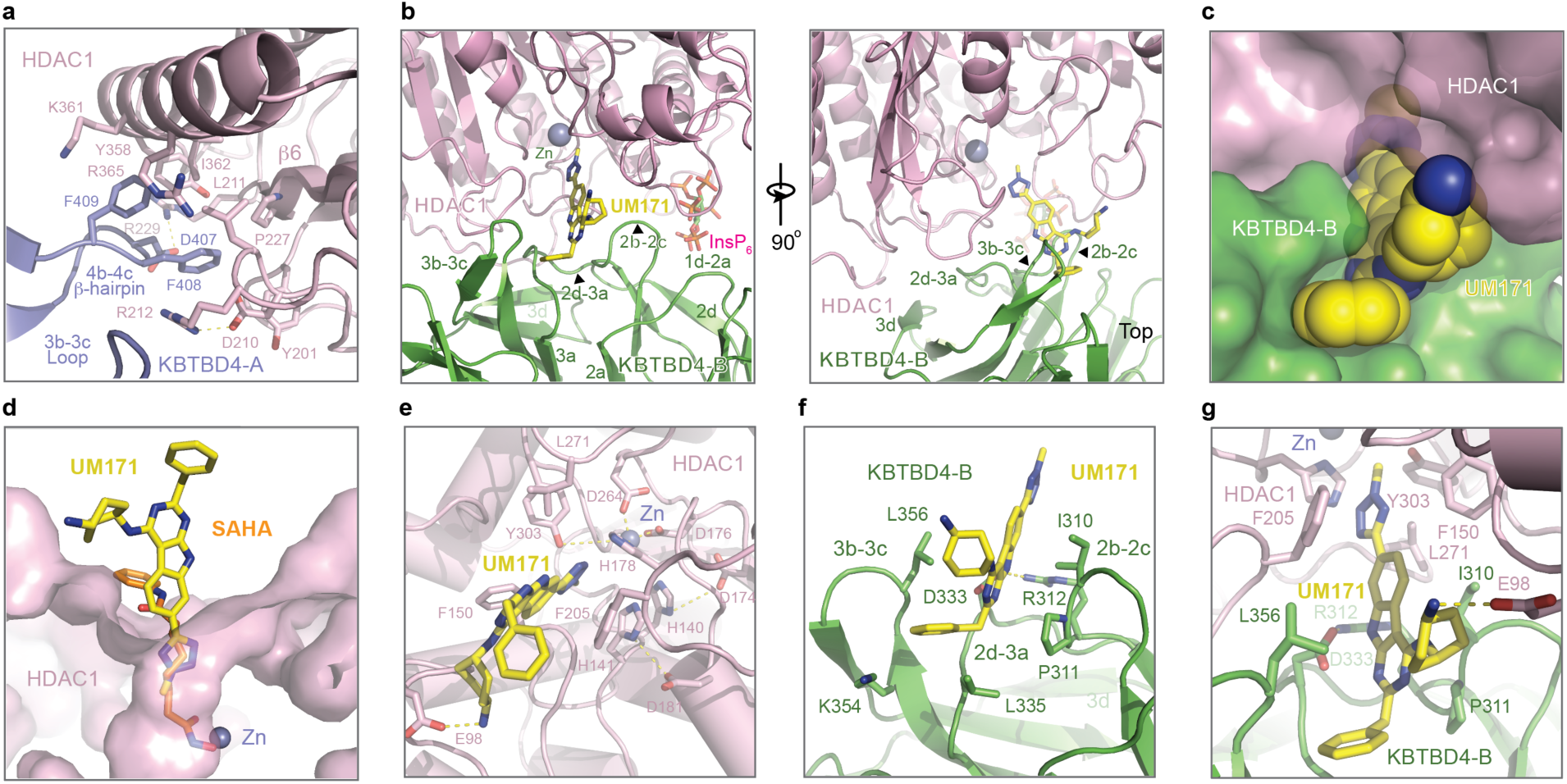
UM171 acts as a molecular glue. **a)** The interface between the 4b-4c β-hairpin of KBTBD4-A (slate) and HDAC1 (pink). The side chains of the interacting amino acids are shown in sticks. **b)** Two orthogonal views of the interface formed between KBTBD4-B (green) and HDAC1 (pink) with UM171 shown in yellow and blue sticks and InsP_6_ shown in green and red sticks. Secondary structures involved in protein-protein interactions are annotated. Zinc (Zn) is shown in sphere (slate). **c)** A close-up view of the surface complementarity among KBTBD4-B (green), HDAC1 (pink), and UM171 (yellow and blue spheres) **d)** A comparison of UM171 (yellow and blue sticks) with SAHA (orange, blue, and red sticks) binding to the active site pocket of HDAC1 (pink surface) with zinc (Zn) shown in slate sphere based on the HDAC2-SAHA structure (PDB: 4LXZ) superimposed with HDAC1. **e)** A close-up view of UM171 (yellow and blue sticks) binding to the active site pocket of HDAC1 (pink). Important residues demarcating the active site of HDAC1 are shown in sticks with zinc (Zn) shown in slate sphere. Potential hydrogen bonds are shown in dashed lines. **f)** A close-up view of UM171(yellow and blue sticks) binding to the surface of KBTBD4-B (green). The side chains of key UM171-contacting residues are shown in sticks. **g)** A close-up view of the UM171 (yellow and blue sticks) binding to the surface pocket formed between KBTBD4-B (green) and HDAC1 (pink). Side chains of select UM171-contacting residues are shown in sticks.

### Interface among HDAC1, KBTBD4-B, and UM171

In cooperation with UM171, the KELCH-repeat domain of KBTBD4-B plays the predominant role in binding HDAC1 by forming an extensive interface with the substrate-binding site of the enzyme (**Fig. 3a**). The HDAC1-binding region of KBTBD4-B encompasses the lateral surface of blade 2 and 3 of its propeller domain and involves multiple secondary structural elements, including the outer d strand of blade 3 (3d) and the 1d-2a, 2b-2c, 2d-3a, and 3b-3c loops (**Fig. 4b**). Comprising both solvent-exposed hydrophobic and polar residues, these KBTBD4-B structural elements present a relatively flat surface and interact with more than half of the active site loops on one side of the deacetylase. In doing so, KBTBD4-B contacts one half of the rim surrounding the active site pocket of HDAC1. In particular, the tip of the KBTBD4-B 2b-2c hairpin loop physically occupies the region of the deacetylase that is involved in recognizing the two amino acids downstream of histone H4 K16, a substrate of HDAC1 (**Extended Data Fig. 6e**)^39^. Despite the spatial proximity of the E3 to the active site pocket of HDAC1, none of the KBTBD4-B residues are positioned close enough to access the catalytic pocket. Such a binding mode creates a suboptimal protein-protein interface with a deep pocket at its periphery, allowing UM171 to insert into and complement this interface (**Fig. 4c, Extended Data Fig. 6f**).

UM171 stabilizes the KBTBD4-B-HDAC1 interactions by bridging both proteins. On the HDAC1 side, UM171 inserts its *N-*methyl-tetrazole moiety into the active site pocket of the enzyme (**Fig. 4d**). The *N*-methyl group of the compound reaches as deep as Cε of the histone H4 Lys16 side chain does (**Extended Data Fig. 6g**). The tetrazole of UM171 is sandwiched between two key residues of HDAC1, Phe150 and Phe205, whose side chains face each other and constrict the active site tunnel leading to the catalytic zinc ion (**Fig. 4e**). The tricyclic pyrimidoindole core lies at the periphery of the pocket, with the cyclohexylamine substituent of UM171 extending outwards to form a salt bridge with a solvent-exposed negatively charged residue, Glu98, of HDAC1. Superposition analysis of UM171-bound HDAC1 and SAHA-bound HDAC2 shows that the corresponding small molecules would competitively occupy the active site, explaining their observed mutual exclusivity (**Fig. 4d**)^40^. However, whereas most HDAC1 active site inhibitors (e.g., SAHA) chelate the catalytic zinc, UM171 does not make direct contact. These findings are consistent with our observations that UM171 cannot directly bind or inhibit HDAC1 alone and instead requires KBTBD4 for LHC engagement and inhibition (**Fig. 2d, e**).

On the KBTBD4-B side, the pyrimidoindole scaffold and the benzyl group of UM171 are tucked into a surface groove between the b-c loops of blade 2 and 3 (**Fig. 4f**). At one end of the groove, the benzyl group of UM171 packs against a hydrophobic wall formed by Pro311, Leu335, and the aliphatic side chain of Lys354. At the other end of the cavity, the pyrimidoindole ring of UM171 is flanked by two hydrophobic residues, Ile310 and Leu356, at the tips of the 2b-2c and 3b-3c loops (**Fig. 4f**). A salt bridge formed between Arg312 and Asp333, which also H-bonds with the UM171 indole N–H, further secures the tricycle. Consistent with these observations, mutation of Pro311 or Asp333 completely blocks CoREST-GFP degradation by UM171 (**Fig. 3e**). In the context of the full complex, Ile310 and Leu356 of KBTBD4-B also make direct hydrophobic interactions with Phe150 and Phe205 of HDAC1, respectively (**Fig. 4g**). These four hydrophobic residues, together with Leu271 of HDAC1, surround UM171 and nucleate a hydrophobic core at the protein-protein interface. Interestingly, the surface groove of the E3 is completely closed on the KBTBD4-A propeller and incompatible with UM171 binding (**Extended Data Fig. 6h**). The b-c loops of blade 2 and 3 in KBTBD4-B, therefore, are most likely spread open by the small molecule with the support of HDAC1. These observations likely explain why UM171 does not show any detectable affinity towards the free KBTBD4 protein and only a single small molecule associates with the dimeric E3 assembly due to the necessary involvement of HDAC1.

### InsP_6_ as a second molecular glue

The HDAC1-MTA1 complex can be stabilized by inositol phosphates, which bind to a surface pocket of the complex situated at the interface between HDAC1 and the corepressor^19,20^. In the KBTBD4-LHC-UM171 complex, a clear density of InsP_6_ is present at the expected binding site between HDAC1 and CoREST (**Fig. 3a, 5a, b**). Remarkably, the InsP_6_ molecule also directly contacts KBTBD4-B, likely acting as a second molecular glue at the KBTBD4-LHC interface. Although we cannot definitively assign the carbon atoms of InsP_6_, all six of its phosphate groups interact with the protein subunits.

**Figure 5.**
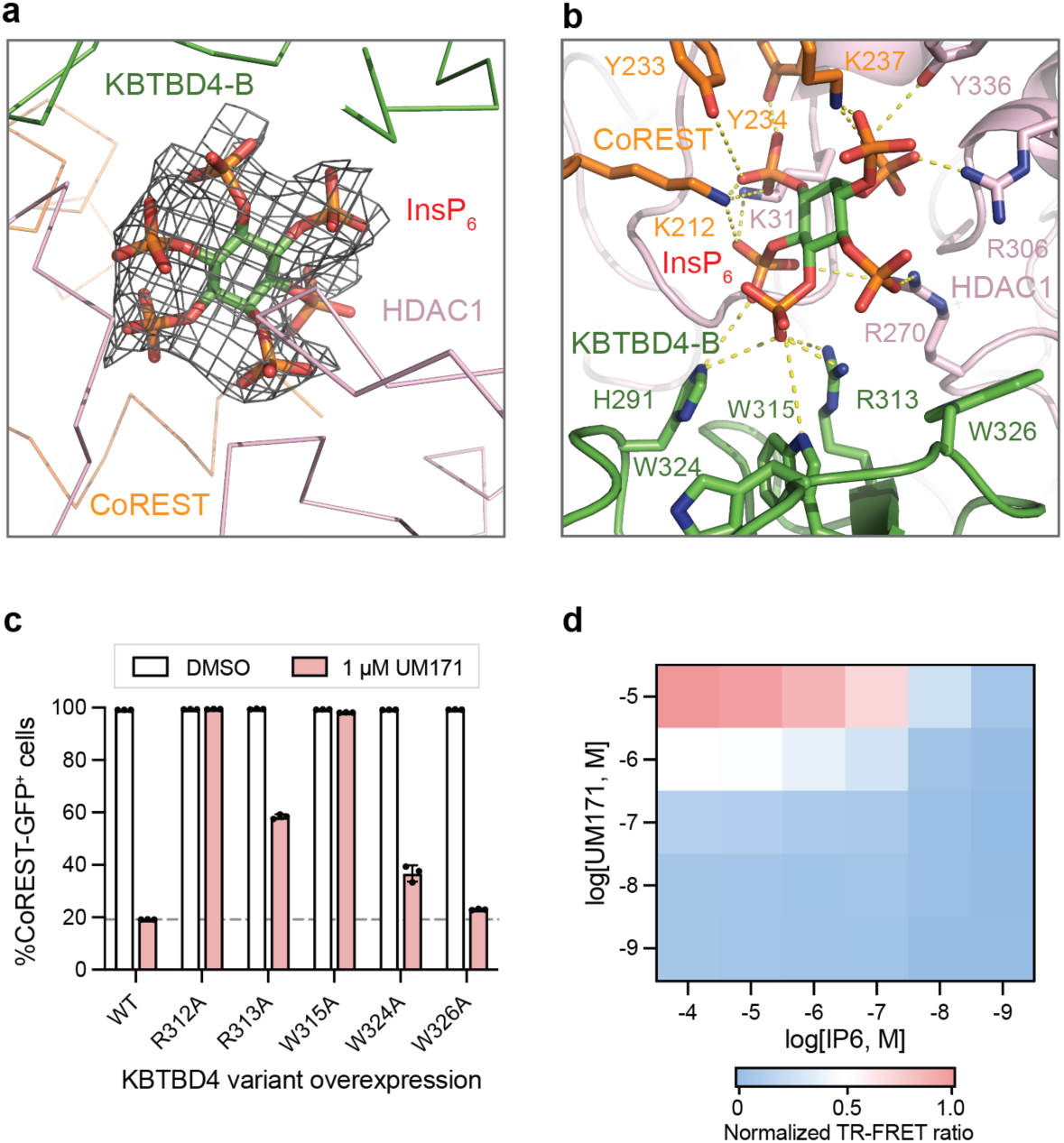
InsP_6_ establishes a bimolecular glue interface. **a)** A close-up view of InsP_6_ (red, orange, and green sticks) with its density map (dark grey colored mesh) at the tri-molecular junction among KBTBD4-B (green), HDAC1 (pink) and CoREST (orange). **b)** A close-up view of the interactions made by InsP6 (red, orange, and green sticks) to the KBTBD4-B (green), HDAC1 (pink) and CoREST (orange) with residues involved in interaction highlighted in sticks. Potential salt bridges and hydrogen bonds are shown in dashed lines. **c)** Flow cytometry quantification of K562 KBTBD4-null CoREST-GFP cells overexpressing indicated KBTBD4 variants after treatment with DMSO or UM171 (1 µM) for 24 h. Bars represent mean ± s.d. of *n = 3* replicates. **d)** Heatmap showing normalized TR-FRET signal between fluorescein-LHC (40 nM) and anti-His CoraFluor-1-labeled antibody with His-KBTBD4 (40 nM) in the presence of increasing concentrations of InsP_6_ (*x*-axis) and UM171 (*y-*axis). Data represent mean of *n = 2* replicates. Results in **c** and **d** are representative of two independent experiments.

On the LHC side, five phosphate groups in InsP_6_ are coordinated by three positively charged residues and a tyrosine residue in HDAC1 (Lys31, Arg270, Arg306, and Tyr336) as well as two positively charged residues and two tyrosine residues from the CoREST SANT1 domain (Lys212, Tyr233, Tyr234, and Lys237). On the KBTBD4-B side, a histidine residue from the 1d-2a loop (His291) and an arginine residue from the 2b-2c loop (Arg 313) each form a salt bridge with one of the six phosphates. InsP_6_ makes an additional contact with Trp315, which is sandwiched between the two basic residues. Interestingly, Trp315 belongs to a cluster of three tryptophan residues on the lateral surface of the KBTBD4-B propeller (Trp315, Trp324, and Trp326), which might be involved in protein-protein interaction. Mutation of these tryptophan residues to alanine revealed that only Trp315 is essential for CoREST degradation by UM171 (**Fig. 5c**).

The binding mode of InsP_6_ in the KBTBD4-LHC complex strongly suggests that the inositol phosphate molecule synergizes with UM171 to further enhance the stability of the E3-neosubstrate complex. Consistent with this notion, UM171 is not sufficient to promote KBTBD4-LHC complex formation in the absence of InsP_6_ and vice versa (**Fig. 2d**). To further explore this mutual dependency, we performed a dose titration of UM171 and InsP_6_ and measured KBTBD4-LHC binding using TR-FRET, revealing that complex formation was only observed in the presence of both small molecules and that they synergistically cooperate (**Fig. 5d**). Altogether, our results reveal the remarkable dependence of a quaternary complex on two small molecule glues in shaping an extensive, induced protein-protein interface.

### Base editing functionally maps the KBTBD4-UM171-HDAC1 interface

To test in situ the interactions identified in the cryo-EM structure, we sought to systematically mutate HDAC1 and KBTBD4 in cells and measure the subsequent impact on CoREST degradation by UM171. Overexpressed HDAC1 could not recapitulate ternary complex formation with UM171 and KBTBD4 (**Extended Data Fig. 7a**) — likely due to non-physiological expression that interferes with endogenous complex stoichiometry. Consequently, we used base editor scanning^41^ to systematically mutate endogenous HDAC1, specifically employing the expanded PAM variant SpG Cas9 cytidine and adenosine base editors (CBE and ABE, respectively) to improve aa mutational coverage (**Fig. 6a**; 428/482 residues for HDAC1 (88.6%), 460/534 residues for KBTBD4 (86.1%))^42^. Due to the redundancy of HDAC1 and HDAC2^26^, we first generated CoREST-GFP knock-in cell lines containing *HDAC2* knockout to circumvent compensation during the base editor scanning (**Extended Data Fig. 7b-d**). Base editors and the pooled sgRNA library targeting *HDAC1* were introduced into K562 CoREST-GFP HDAC2-null cells by lentiviral transduction. After puromycin selection, transduced cells were treated with UM171 (24 h, 1 µM) or vehicle control, and cells remaining GFP positive were FACS-sorted. Enriched sgRNAs were identified to reveal HDAC1 positions required for CoREST-GFP degradation (i.e., greater sgRNA enrichment scores) (**Fig. 6b**). As expected, sgRNAs predicted to introduce missense mutations were most highly enriched in the base editor scanning, suggesting that they may confer resistance to UM171-induced CoREST degradation (**Extended Data Fig. 7e**). Due to the non-uniform coverage of sgRNAs, we next used LOESS regression to estimate per-residue enrichment scores from the measured sgRNA scores and then compared them to a simulated distribution generated by shuffling sgRNA scores. This allowed us to assign a linear clustering score in order to determine if a given residue may be more enriched than expected by chance (**Extended Data Fig. 7f**, see Methods)^43^. Generally, this method assigns greater significance to short intervals along the linear coding sequence that contain multiple enriched sgRNA hits.

**Figure 6.**
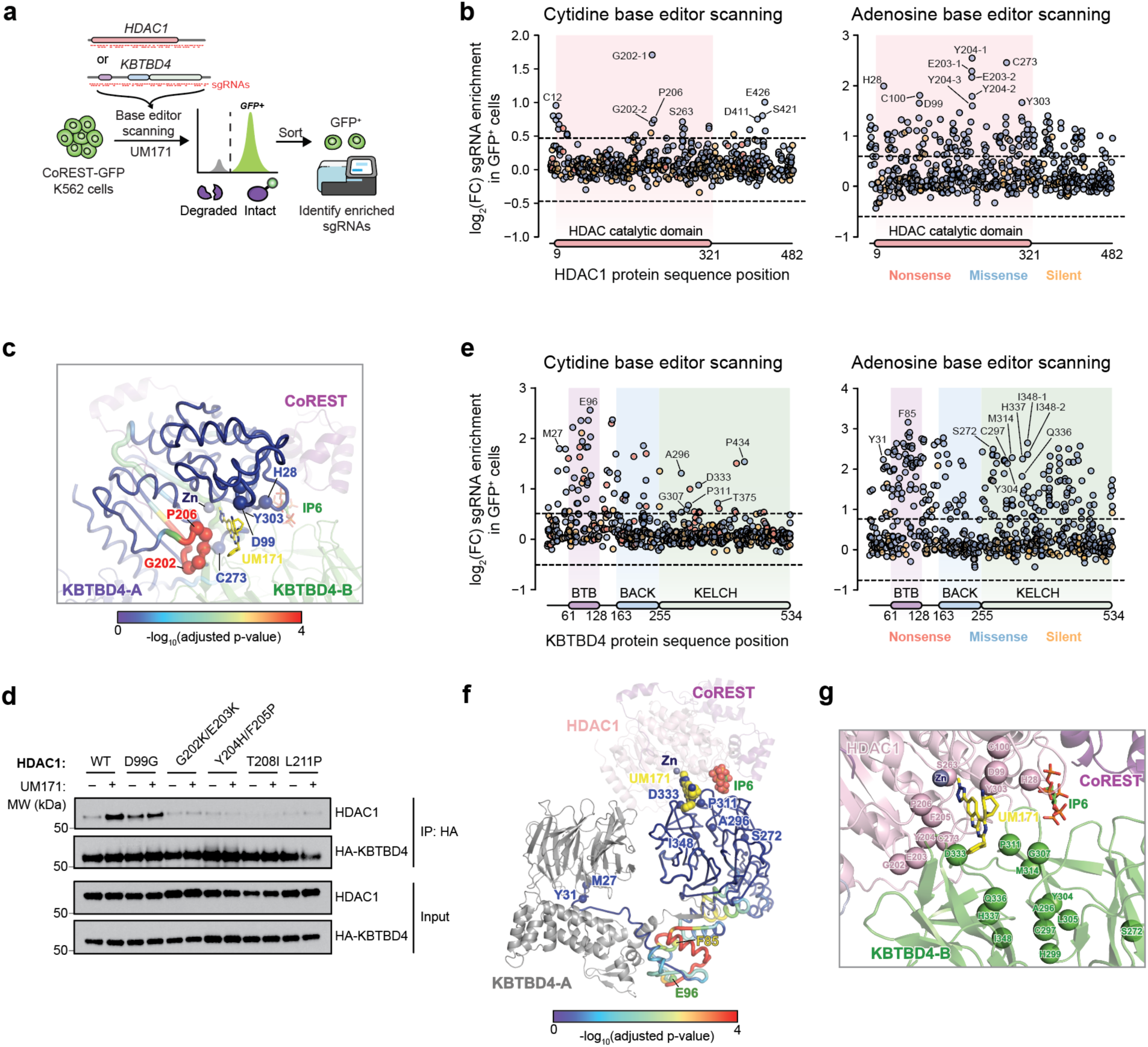
Base editing functionally maps the KBTBD4-UM171-HDAC1/2 interface. **a)** Schematic of base editor scanning of *HDAC1* and *KBTBD4* in K562 CoREST-GFP cells. **b)** Scatter plots showing log_2_(fold-change sgRNA enrichment) (*y-*axis) in GFP^+^ cells versus unsorted cells for base editor scanning of *HDAC1* using cytidine base editor (left) and adenosine base editor (right). Dotted lines represent ± 4 s.d. from the mean of non-targeting controls (n = 199). Selected sgRNA hit positions are labeled. **c)** Structure of HDAC1-CoREST-KBTBD4 showing HDAC1 residues colored by linear clustering score from base editor scanning (see **Extended Data Fig. 7f**). Cα-positions of selected top-enriched sgRNAs, marked in **Fig. 6b**, are shown as spheres. **d)** Immunoblots of HA-KBTBD4 IP in the presence of DMSO or 1 µM UM171 (1 h), and 1 µM MLN4924 (3 h) from clonal 293T cell lines, containing indicated HDAC1 base edits, transfected with HA-KBTBD4 (*n* = 2). See **Extended Data Fig. 8a** for base editing genotypes. **e)** Scatter plots showing log_2_(fold-change sgRNA enrichment) (*y-*axis) in GFP^+^ cells versus unsorted cells for base editor scanning of KBTBD4 using cytidine base editor (left) and adenosine base editor (right). Dotted lines represent ± 4 s.d. from the mean of non-targeting controls (n = 199). Selected sgRNA hit positions are labeled. **f)** Structure of HDAC1-CoREST-KBTBD4 showing KBTBD4-B residues colored by linear clustering score from base editor scanning (see **Extended Data Fig. 7f**). Cα-positions of selected top-enriched sgRNAs, marked in **Fig. 6e**, are shown as spheres. **g)** View of the HDAC1-UM171-KBTBD4 interface showing Cα-positions of selected top-enriched sgRNAs, marked in **Fig. 6e, 6f**, as spheres. Results in **d** are representative of two independent experiments.

Mapping these per-residue linear clustering scores for HDAC1 onto our cryo-EM structure revealed a strong mutational hotspot surrounding the UM171 binding site at the HDAC1-KBTBD4-B interface (**Fig. 6c**). In particular, this included a loop comprising HDAC1 Gly202–Pro206. When considering individual enriched sgRNAs, many of the corresponding base edits were predicted to alter residues that directly contact UM171 as well as KBTBD4-B (e.g., sgH28, sgD99, sgG202, sgE203, sgY204, sgP206, sgS263, sgC273, sgY303), suggesting that they disrupt multiple aspects of complex formation. To validate these base edits, we generated and genotyped clonal 293T cell lines in which HDAC1 was edited by sgD99, sgG202-2, or sgY204-1 (**Extended Data Fig. 8a**). We also generated cell lines using sgRNAs targeting residues proximal to the KBTBD4-A contact site (sgT208, sgL211). We confirmed that these base edits introduced into HDAC1 were sufficient to block KBTBD4-HDAC1 co-IP promoted by UM171 (**Fig. 6d**), validating the hotspot identified by base editor scanning and showing that this interface is critical for the E3-neo-substrate complex formation.

Using an analogous base editing strategy, we next identified functional residues on KBTBD4 (**Fig. 6e**). We validated that a subset of CBE base edits block CoREST-GFP degradation by individual sgRNA transduction (**Extended Data Fig. 8b**). As anticipated, many more sgRNAs targeting KBTBD4 scored as enriched hits, as any significant loss-of-function mutation in the E3 can block degradation or even act in a dominant negative manner, in contrast to mutations in the neo-substrate (i.e., HDAC1). Most top-enriched sgRNAs (CBE: 18/80; ABE: 47/197) targeted the BTB domain, and the corresponding base edits likely disrupt KBTBD4’s homodimerization and/or interaction with CULLIN3 and hence ligase activity. In line with this notion, linear clustering analysis of the KBTBD4 base editor scanning data showed that the strongest mutational hotspot resided in the BTB domain and along the dimerization interface (**Fig. 6f; Extended Data Fig. 7f**), supporting the idea that dimerization is critical for KBTBD4 function. Notably, several top-enriched base edits targeted residues within or contacting the N-terminal β1 strand, further validating its importance in KBTBD4 function.

Many top-enriched base edits are also predicted to alter various blades of the KELCH domain. Regions containing these individual top-enriched sgRNAs did not score highly in the linear clustering, likely due to the overwhelmingly strong hits in the dimerization interface as well as the 3D structure of the β-propeller domain. In particular, several top-enriched base edits target regions surrounding the UM171 binding site in-between blade 2 (2b-2c loop) and blade 3 (2d-3a loop) (**Fig. 6f, g**). We further investigated the base edits produced by sgA296, sgP311, and sgD333, first confirming the predicted base editing outcomes by genotyping and then the stability of the corresponding KBTBD4 variants in cells (**Extended Data Fig. 8c, d**). Overexpression of these KBTBD4 variants (i.e., A296T, P311L, D333N) in K562 KBTBD4-null CoREST-GFP cells showed that CoREST-GFP remained stable in the presence of UM171 (**Fig. 3e**), confirming that these variants likely disrupt UM171 binding. Taken together, base editor scanning of the neo-substrate and E3 support, in the native context, the complex interfaces defined by our LHC-UM171-KBTBD4 cryo-EM structure.

## Discussion

Targeted protein degradation is a powerful strategy for drug discovery. While rational approaches to identifying degraders have been intensely pursued, most molecular glue degraders have been identified serendipitously^4,5^. Deconvoluting the mechanism of action of these small molecules, nevertheless, has transformed the field of targeted protein degradation, unveiling new strategies and E3 ligase scaffolds for proximity-induced pharmacology and drug discovery. In this study, we elucidate the molecular and structural mechanisms of UM171, establishing its function as a molecular glue. By engaging with the active site of HDAC1, UM171 facilitates the formation of a protein-protein complex involving LHC, InsP_6_, and an asymmetric dimeric KBTBD4 complex. Our cryo-EM structure is functionally substantiated by our base editor scanning experiments, showcasing the synergy of these approaches. The sensitivity of molecular glues to endogenous protein stoichiometry highlights a powerful advantage of using base editing to map small molecule interaction sites.

The cryo-EM structure of the LHC-UM171-InsP_6_-KBTBD4 complex provides the first example of a molecular glue-licensed CRL3 E3 engaged with its neo-substrate. Among the superfamily of CRL E3s, CRL3 has the most substrate receptors, the majority of which share the same BTB-KELCH domain composition as KBTBD4^44–48^. Our results, therefore, substantially expand the repertoire of human E3 ligases that are potentially rewireable by molecular glue degraders. Importantly, in contrast to most CRL4s, CRL3s and some members of other CRL families are known to function as homodimers with two substrate-binding domains, allowing cooperative binding of two degrons encoded within a single substrate polypeptide. Rather than co-opting each substrate-binding domain individually, a single molecule of UM171 leverages both protomers of the E3 to engage HDAC1 cooperatively and asymmetrically as a single assembly. Such an E3-glue-neo-substrate architecture has not been observed before and underscores the unique potentials of CRL3s and other dimeric CRLs to be reprogrammed by small molecules.

Remarkably, UM171 binds a pocket that is only present upon KBTBD4-B-HDAC1 contact, fitting a common mechanistic theme in which molecular glues stabilize complementary yet sub-optimal interaction surfaces between two proteins^5,6^. Interestingly, unlike the HDAC1 active site, the KBTBD4-B surface groove occupied by UM171 is only induced upon E3-neo-substrate complex formation, allowing a single UM171 molecule to asymmetrically engage a dimeric E3 complex. This implicates that the surface of E3 scaffolds can be structurally plastic and might contain more binding sites for molecular glue engagement than their apo structures suggest. We also demonstrate the requirement of InsP_6_ as a second molecular glue at this protein-protein interface, which indicates that UM171 is insufficient to induce stable productive interaction between KBTBD4 and HDAC1. With the assistance of a cellular metabolite, a synthetic compound, therefore, can function as an efficacious molecular glue degrader, despite its inability to do so independently.

We establish HDAC1/2 as the unexpected, elusive target of UM171, highlighting novel modalities to target this enzyme class. Notably, UM171 does not inhibit HDAC1/2 activity alone and only leads to the potent degradation of select HDAC1/2 corepressors, despite KBTBD4 making no direct contact with the corepressor itself. This is possibly due to the presence of neighboring domains in undegraded corepressors (i.e., BAH2 domain in MTA1) that crowd the HDAC1 active site or surrounding areas, sterically blocking KBTBD4 engagement^49^. By leveraging the KBTBD4 E3 ligase, UM171 demonstrates an unprecedented ability to preferentially down-regulate selective subunits of HDAC1/2 complexes (i.e., CoREST, MIER1), which are otherwise not readily amenable to therapeutic targeting. Lastly, the mechanism of UM171 provides another compelling example of how active site ligands can serendipitously act as molecular glues, further highlighting that enzyme active sites may be privileged as partners in pharmacologically-induced protein-protein interactions^50^. In summary, this work reveals the remarkable molecular sophistication of molecular glue degraders in reprogramming extensive protein-protein contacts and the vast opportunities for their prospective discovery.

## Acknowledgements.

The authors would like to thank J. Nelson at the Bauer Core Facility of Harvard University for assistance with FACS and J.D. Quispe and S. Dickinson at the Arnold and Mabel Beckman Cryo-EM Center of the University of Washington, R. Yan, X. Zhao, J. Jung, and Z. Yu at the Cryo-EM Facility on the Janelia Research Campus of the Howard Hughes Medical Institute, and T. Humphreys and M. Campbell at Fred Hutch EM & cryo-EM Core for their assistance in electron microscopy data acquisition, as well as D. Asarnow from the Veesler laboratory at the University of Washington for his technical insights and suggestions, and members of the Zheng and Liau laboratories, especially D.V. Rusnac, S. Zhang, H. Shi, E. Garcia, J. Woods, and N. Lue, for their discussion and inputs. The authors would also like to thank E. Svenningsen, L. Zhi, and C. Woo for assistance with target identification experiments. A.M. is supported by PRIN2020 from the Italian Ministry for Research MUR. K.L. is supported by the American Heart Association (Postdoctoral Fellowship Award 826614). N.C.P is supported by the National Science Foundation (DGE1745303). P.A.C. is supported by the National Institute of General Medical Sciences (R35GM149229) and the Leukemia and Lymphoma Society. B.L. is supported by the Ono Pharma Foundation, the Camille and Henry Dreyfus Foundation (Teacher-Scholar Award), the Blavatnik Accelerator Fund (Harvard University), the National Institute of General Medical Sciences (1DP2GM137494), and the National Cancer Institute (1R01CA274437). N.Z. is supported by the Howard Hughes Medical Institute.

## Author contributions

B.L. and N.Z. conceived the project with inputs from M.Y., O.Z., X.X., and P.M.G.. M.Y. and P.M.G. performed cellular experiments, flow cytometry, and base editor scanning. O.Z. purified KBTBD4 for biochemical studies and conducted FP assays. O.Z. and H.J. conducted in vitro ubiquitination experiments, O.Z., E.N., and K.L. purified LHC for biochemical and structural studies, and Z.W. conducted LHC deacetylation assays, with input from P.A.C.. O.Z. and N.C.P. conducted TR-FRET experiments and HDAC activity assays with input from R.M.. C.L. and M.Y. conducted computational analysis of base editor scanning. J.L., N.C., S.H., and I.B. synthesized UM171 and derivatives. O.Z., H.S.K., I.I., and A.L.W. assisted with cellular experiments. N.C. and M.T. conducted proteomics with input from L.B-P.. M.B. performed biochemical studies, with input from A.M.. H.M. and X.X. prepared CUL3-RBX1 for in vitro ubiquitination assay. H.M. and X.X. purified KBTBD4 for structural studies. X.X. performed sample preparation, cryo-EM grid preparation, specimen screening, data collection and processing. X.X., N.Z., and B.L. analyzed the structures. N.Z. and B.L. held overall responsibility for the study.

## Supplementary Materials

**Extended Data Figure 1.**
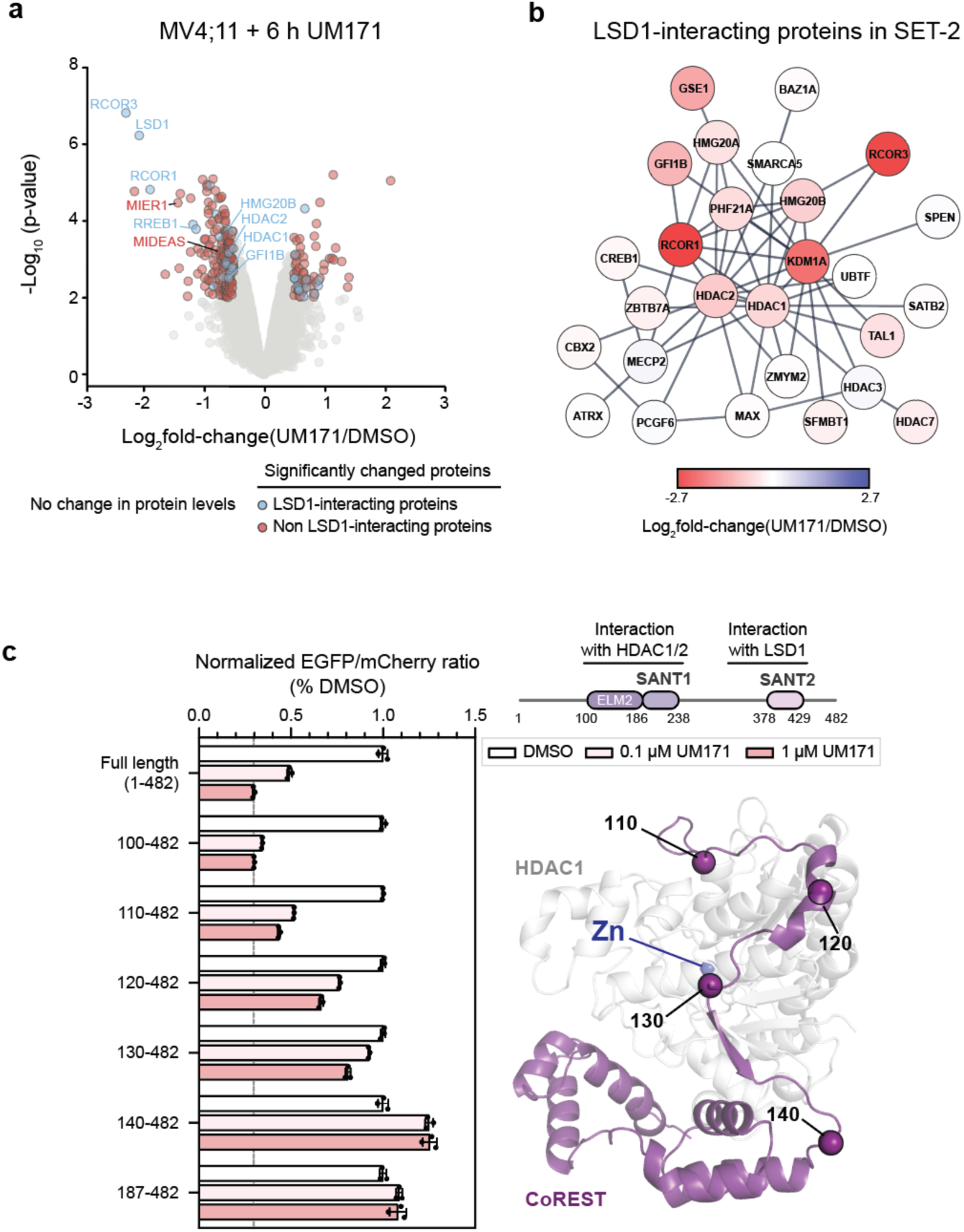
Supporting Data for Figure 1. **a)** Volcano plot showing proteins up- and down-regulated upon UM171 treatment in MV4;11 cells. Colored dots (blue, red) show proteins with | log_2_(fold-change) | > 0.5 in UM171 versus DMSO treatment and p-value < 0.01. Dots in red and blue depict proteins enriched or absent in LSD1 co-IP/MS, respectively. **b)** Protein STRING network showing proteins enriched in LSD1 co-IP/MS in SET-2 cells. Color scale depicts log_2_(fold-change) in UM171 versus DMSO treatment. **c)** Left: GFP to mCherry fluorescence ratio measured by flow cytometry quantification of MOLM-13 cells expressing the indicated CoREST-GFP reporter treated with DMSO or UM171 (0.1 or 1 µM) for 24 h. Bars represent mean ± s.d. of *n =* 3 replicates. Right: Pymol alpha-fold structure of HDAC1-CoREST with key positions and Zn highlighted. Results in **a** and **b** are representative of one independent experiment conducted in technical triplicates. Results in **c** are representative of two independent experiments.

**Extended Data Figure 2.**
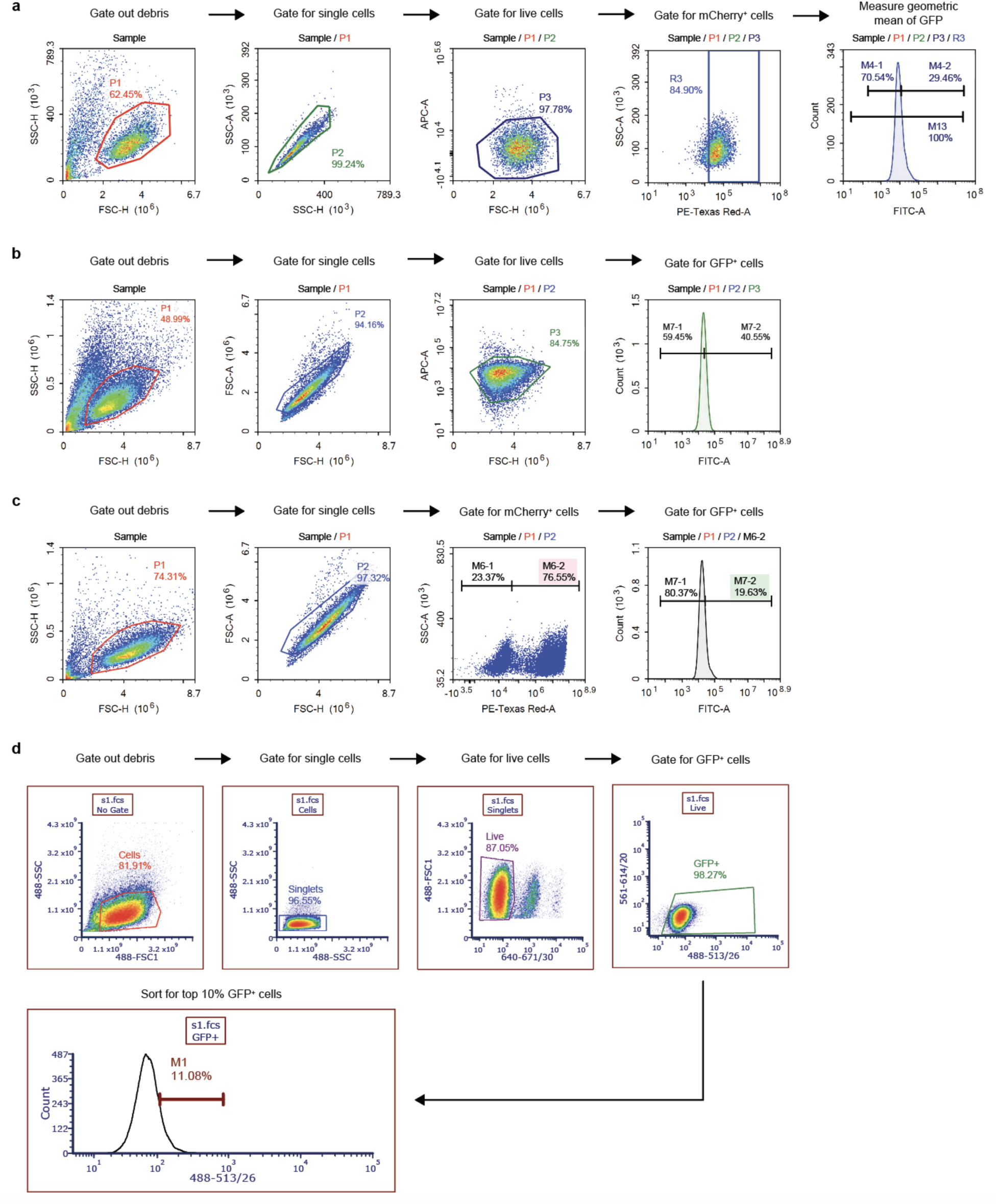
Representative Gating Schemes for Flow Cytometry. **a)** Representative gating scheme for flow cytometric analysis of fluorescence-based corepressor degradation assays in MOLM-13 cells. Helix NP NIR was used as a viability dye. GFP fluorescence and mCherry fluorescence were monitored on the FITC and PE-Texas Red channels, respectively. **b)** Representative gating scheme for flow cytometric analysis of CoREST-GFP degradation by KBTBD4. Helix NP NIR was used as a viability dye. GFP fluorescence was monitored on the FITC channels. **c)** Representative gating scheme for flow cytometric analysis of CoREST-GFP degradation by KBTBD4 overexpression in KBTBD4-null K562 cells. GFP fluorescence and mCherry fluorescence were monitored on the FITC and PE-Texas Red channels, respectively. **d)** Representative gating scheme for FACS-based sorting of base editor screens in CoREST-GFP K562 cells. Helix NP NIR was used as a viability dye. GFP fluorescence was monitored on the FITC channels.

**Extended Data Figure 3.**
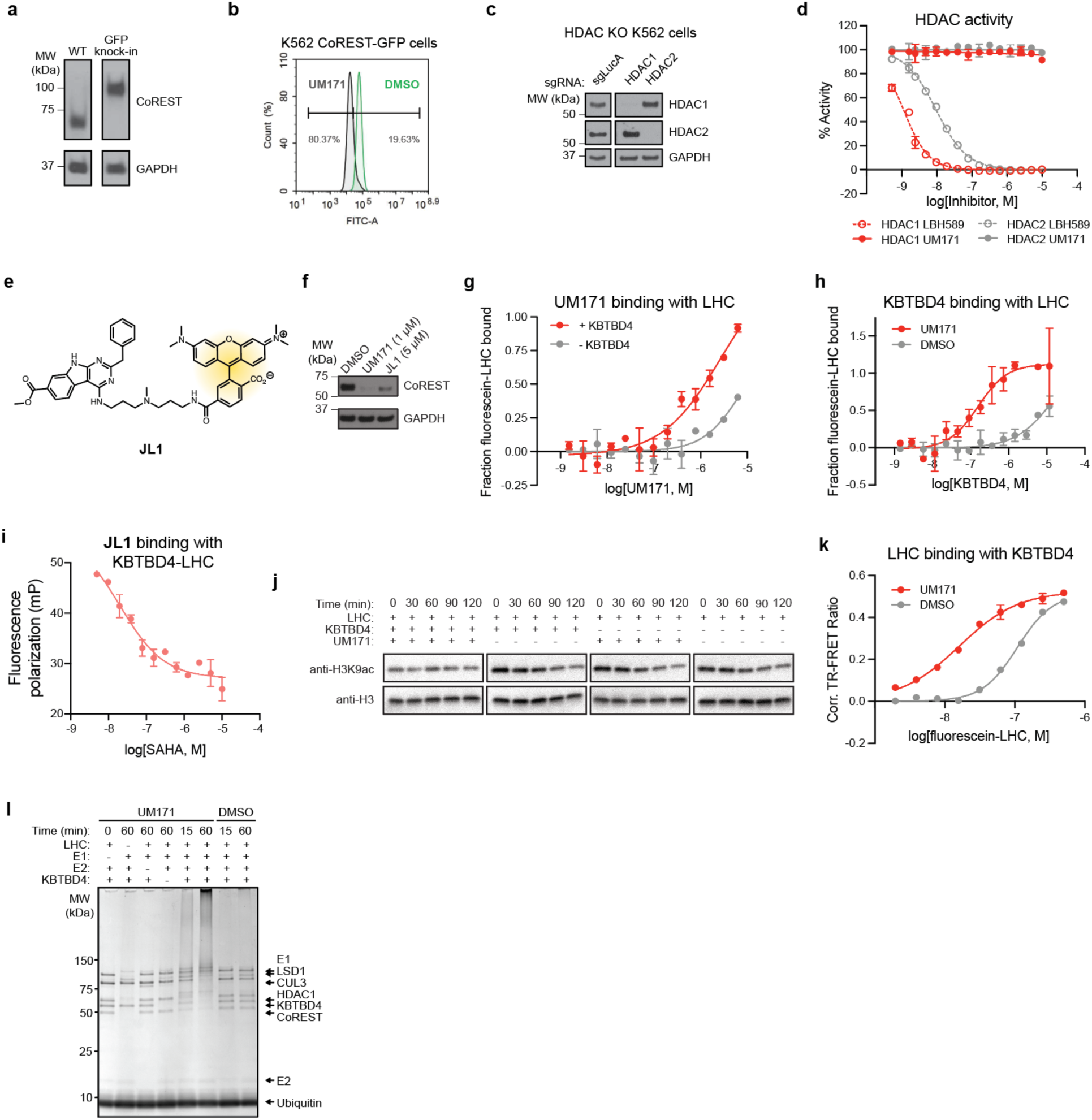
Supporting Data for Figure 2. **a)** Immunoblot showing CoREST and GAPDH for wild-type and CoREST-GFP K562 cells. **b)** Representative flow cytometry quantification showing GFP signal in K562 CoREST-GFP cells for treatment with DMSO (green) or 1 µM UM171 (gray) treatment for 6 h. Gate was determined where 1% of the DMSO condition are considered FITC^−^. **c)** Immunoblot showing HDAC1, HDAC2, and GAPDH in K562 cells after transduction of the indicated sgRNAs. **d)** Dose-response curves showing relative enzyme activity (*y*-axis) for indicated HDAC in the presence of increasing concentrations of the indicated compounds (*x*-axis). Data represent mean ± s.d. of *n* = 2 replicates. **e)** Chemical structure of JL1. **f)** Immunoblot showing CoREST and GAPDH in K562 cells after treatment with UM171 or JL1 at indicated concentrations. **g)** Binding curves showing fraction of fluorescein-LHC bound (*y*-axis) upon increasing concentrations of UM171 (*x*-axis) in the presence and absence of KBTBD4 (200 nM) with fluorescein-LHC (200 nM). Data represent mean ± s.d. of *n* = 2 replicates. **h)** Binding curves showing fraction of fluorescein-LHC bound (*y*-axis) upon increasing concentrations of KBTBD4 (*x*-axis) in the presence and absence of UM171 (50 µM) with fluorescein-LHC (200 nM). Data represent mean ± s.d. of *n* = 2 replicates. **i)** Dose-response curves showing fluorescence polarization (*y*-axis) of JL1 (10 nM) with KBTBD4 (5 µM) and LHC (20 nM) in the presence of increasing concentrations of SAHA (*x*-axis). Data represent mean ± s.d. of *n* = 2 replicates. **j)** Immunoblot for in vitro deacetylation assays of LHC with H3K9ac modified mononucleosomes staining with indicated antibodies against H3K9ac and total H3 under the indicated conditions. **k)** Dose-response curve showing TR-FRET signal (*y*-axis) between anti-His CoraFluor-1-labeled antibody with His-KBTBD4 (10 nM) and varying concentrations of fluorescein-LHC (*x*-axis) in the presence of DMSO or UM171 (10 µM). Data are mean ± s.d. of *n* = 2 replicates. **l)** Coomassie staining for in vitro ubiquitination assays of CRL3^KBTBD4^ (500 nM) with fluorescein-LHC (500 nM) in the presence of DMSO or UM171 (10 µM). Results in **a-f, j**, and **k** are representative of two independent experiments. Results in **g-I** are representative of one experiment. Results in **l** are representative of three independent experiments.

**Extended Data Figure 4.**
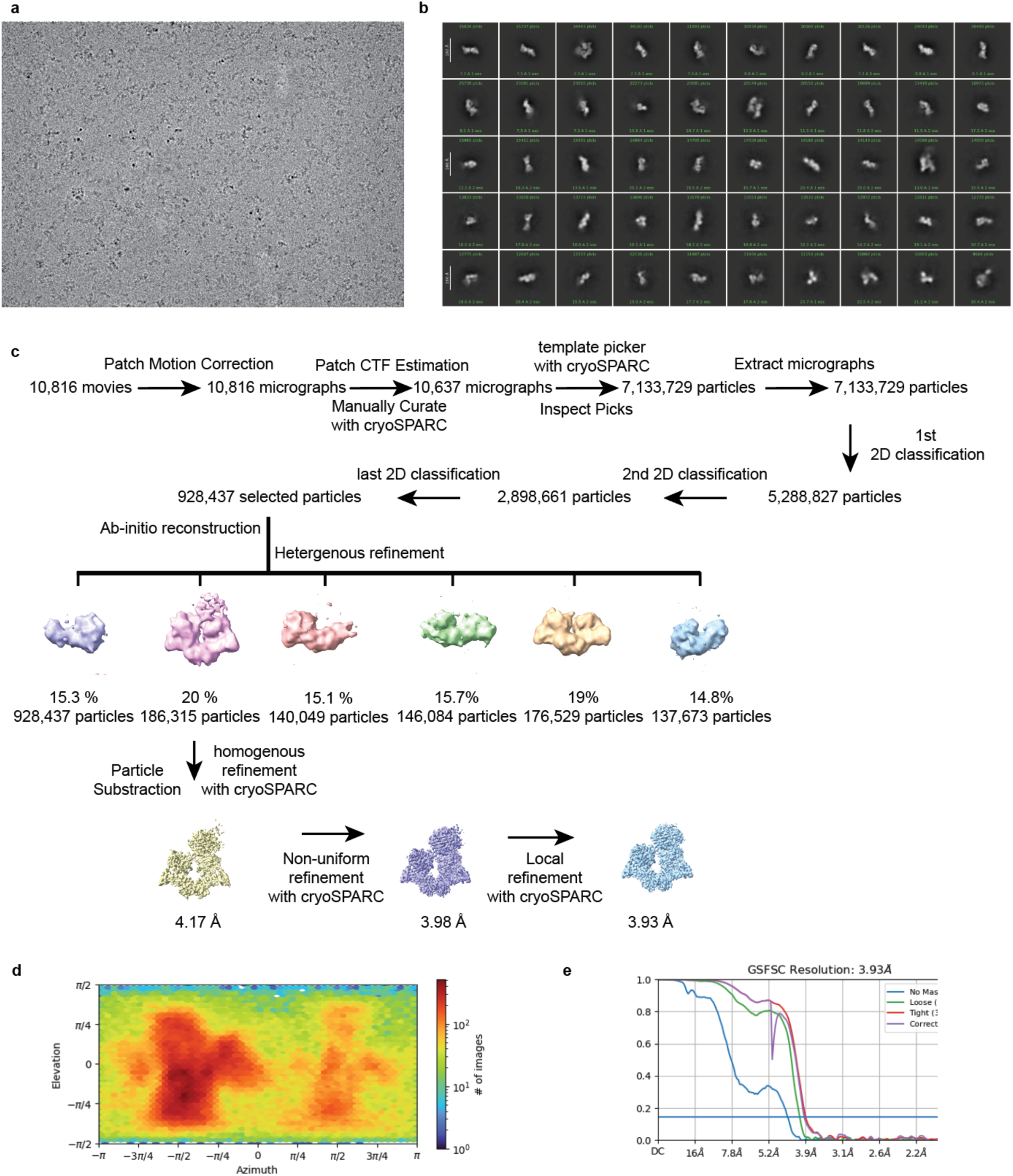
Cryo-EM data processing. **a)** A representative cryo-EM micrograph. **b)** Typical 2D averages of the cryo-EM dataset; scale bar 10 nm. **c)** The flowchart of single particle analysis of the LHC-UM171-KBTBD4 complex. **d)** The angular distribution of particles used in the final reconstruction. **e)** Fourier shell correlation (FSC) curves for LHC-UM171-KBTBD4. At the Gold-standard threshold of 0.143, the resolution is 3.93 Å.

**Extended Data Figure 5.**
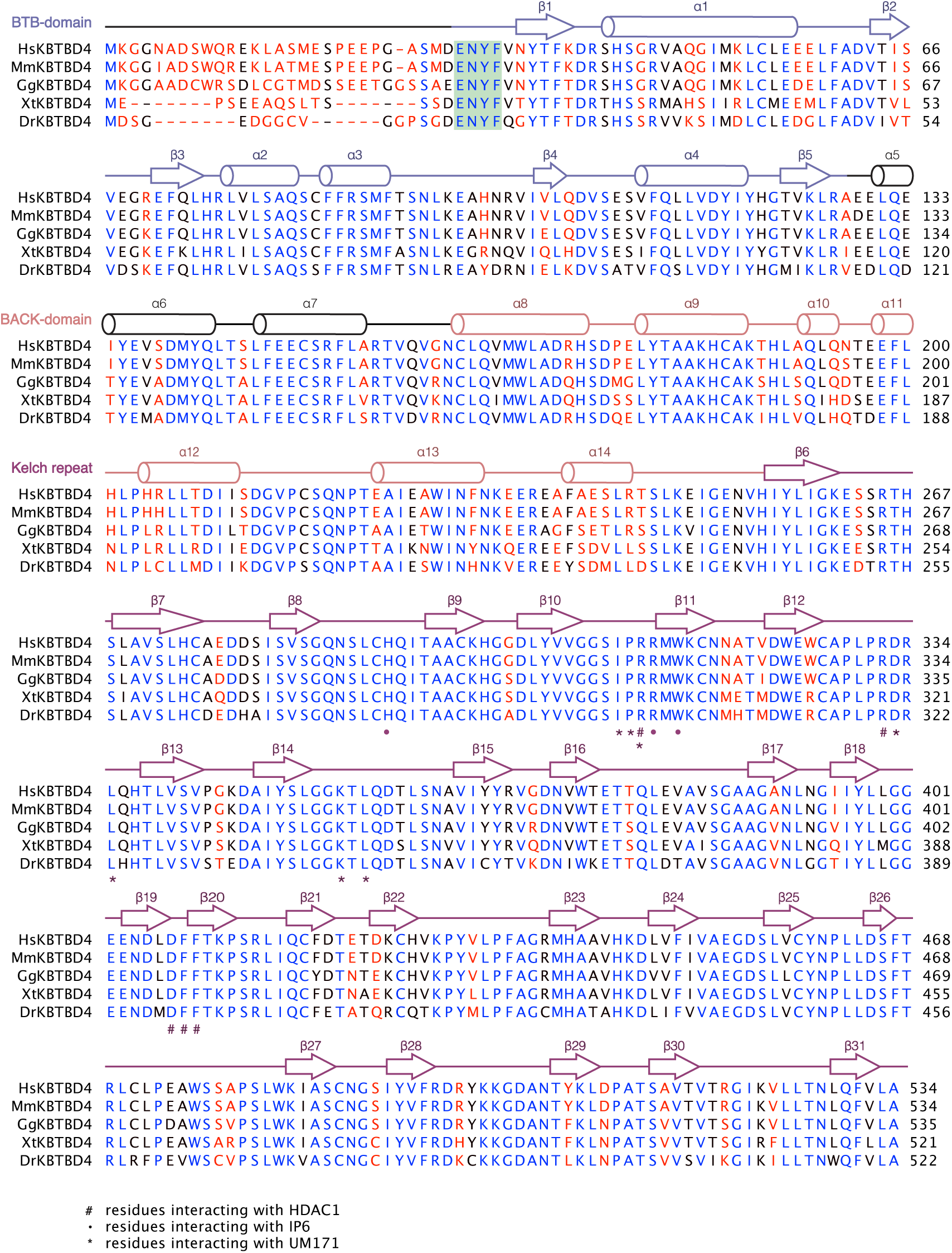
Sequence alignment and structural annotation of KBTBD4. Sequence alignment of five vertebrate KBTBD4 orthologues (Hs: *Homo sapiens*, Mm: *Mus musculus*, Gg: *Gallus gallus*, Xt: *Xenopus tropicalis*, Dr: *Danio rerio*) with second structure annotations. The sequences of the BTB, BACK and KELCH-repeat domains are underlined in different colors (slate, salmon, and purple). The residues interacting with HDAC1 are labeled with “#”; the residues interacting with InsP_6_ are labeled with “•”; and the residues interacting with UM171 are labeled with “*”. Strictly conserved residues are colored in blue.

**Extended Data Figure 6.**
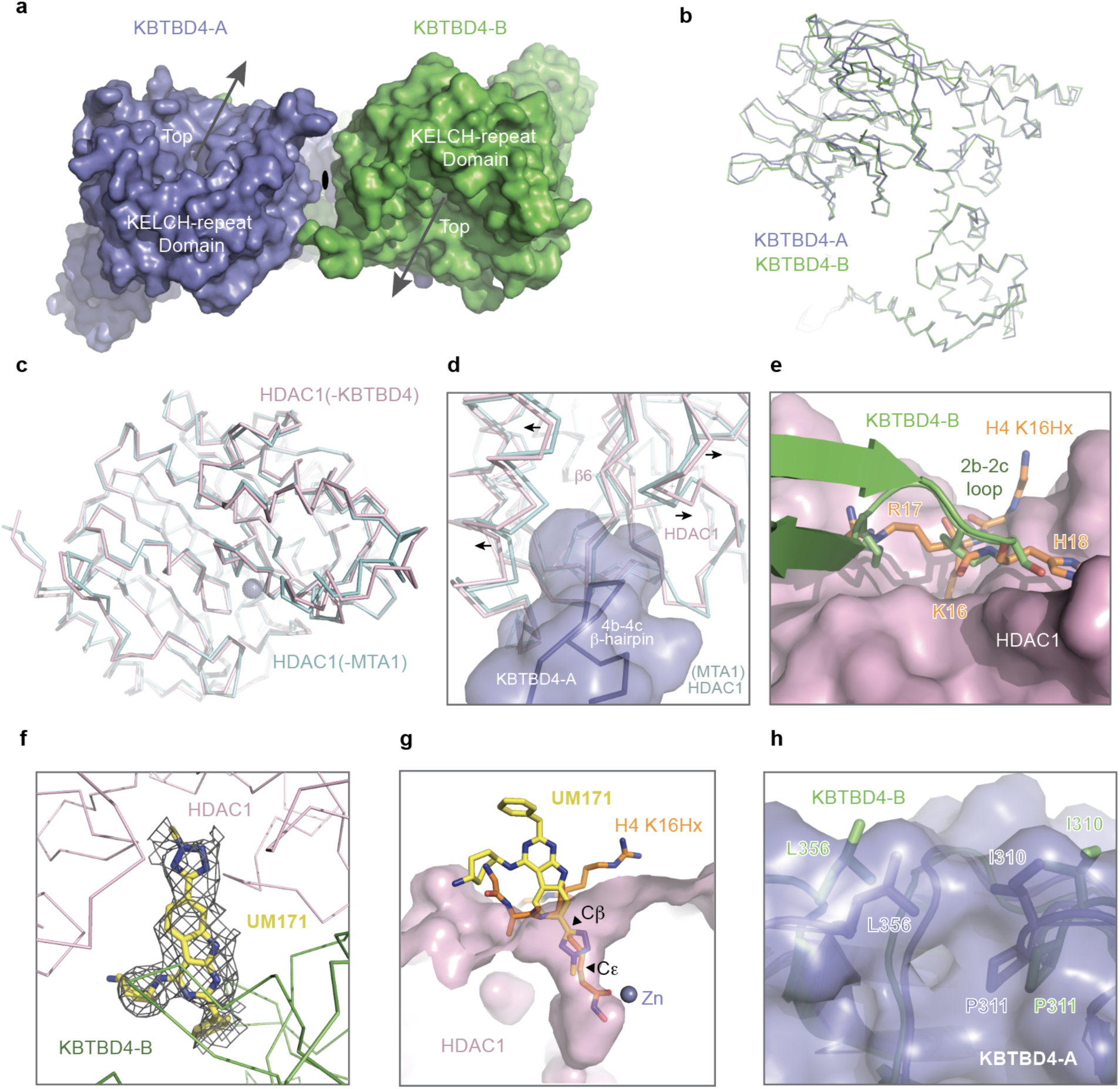
Structural analysis of the KBTBD4-LHC-UM171 complex. **a)** The top view of the KBTBD4 dimer with chain A colored in blue and chain B colored in green in pseudo-two-fold symmetry. **b)** Superposition of KBTBD4 chain A (blue) with chain B (green). **c)** Superposition of HDAC1 bound to KBTBD4 (pink) versus bound to MTA1 (cyan, PDB: 4BKX). **d)** A close-up view of the HDAC1 (pink) region remodeled by the 4b-4c β-hairpin (blue) of the KBTBD4-A (blue surface) versus HDAC1 (cyan) bound to MTA1 (PDB: 4BKX). **e)** A superposition comparison of the 2b-2c loop of KBTBD4-B (green) occupying the active site of HDAC1 (pink) with the histone H4 K16Hx peptide (orange sticks) (PDB: 5ICN). **f)** A close-up view of the UM171 (yellow sticks) along with its density (dark grey mesh) at the interface between HDAC1(pink) and KBTBD4-B (green). **g)** A superposition comparison of the HDAC1-binding modes between UM171 and the histone H4 K16Hx peptide (PDB:5ICN). **h)** A superposition comparison of the UM171 binding region in KBTBD4-B (green) and the corresponding region in KBTBD4-A (blue surface). Residues involved in UM171 binding in KBTBD4-B and their corresponding residues in KBTBD4-A are highlighted in sticks.

**Extended Data Figure 7.**
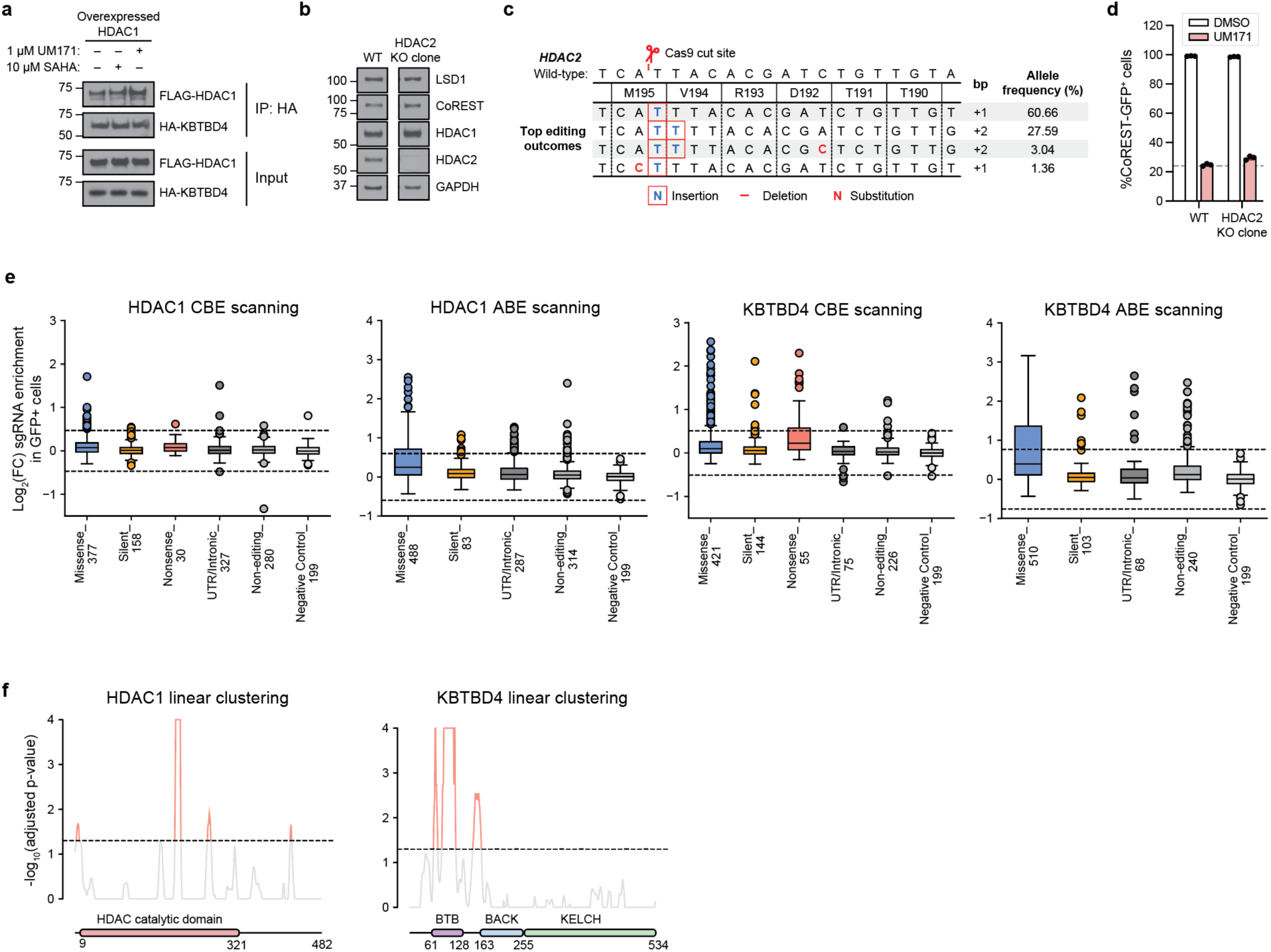
Base editor scanning of HDAC1 and KBTBD4. **a)** Immunoblot of HA IP in the presence of 1 µM UM171, 10 µM SAHA, or vehicle (1 h treatment) from 293T cells transfected with FLAG-HDAC1 and HA-KBTBD4. **b)** Immunoblot showing CoREST-GFP, HDAC1, and HDAC2 in K562 HDAC2-null CoREST-GFP cells. **c)** Allele frequency table for K562 HDAC2-null clonal cell lines. Genotyping was performed once *(n = 1)*. **d)** Flow cytometry quantification showing GFP signal in K562 HDAC2-null CoREST-GFP cells for treatment with DMSO (white) or 1 µM UM171 (pink) treatment for 24 h. **e)** Boxplot of log_2_(fold-change sgRNA enrichment) for HDAC1 and KBTBD4 cytidine base editor (CBE) and adenosine base editor (ABE) scanning. sgRNAs classified by predicted editing outcome and the number of sgRNAs are indicated. Outliers are shown individually. UTR, untranslated region. **f)** Line plots showing −log_10_-transformed (adjusted p*-*values, *P*) for the observed per-residue sgRNA enrichment scores (*y*-axis) plotted against the (left) *HDAC1* CDS (*x*-axis) and (right) *KBTBD4* CDS (*x-*axis). The dotted line corresponds to *P* = 0.05 and residues with *P* ≤ 0.05 are highlighted in red. Results in **a, b**, and **d** are representative of two independent experiments.

**Extended Data Figure 8.**
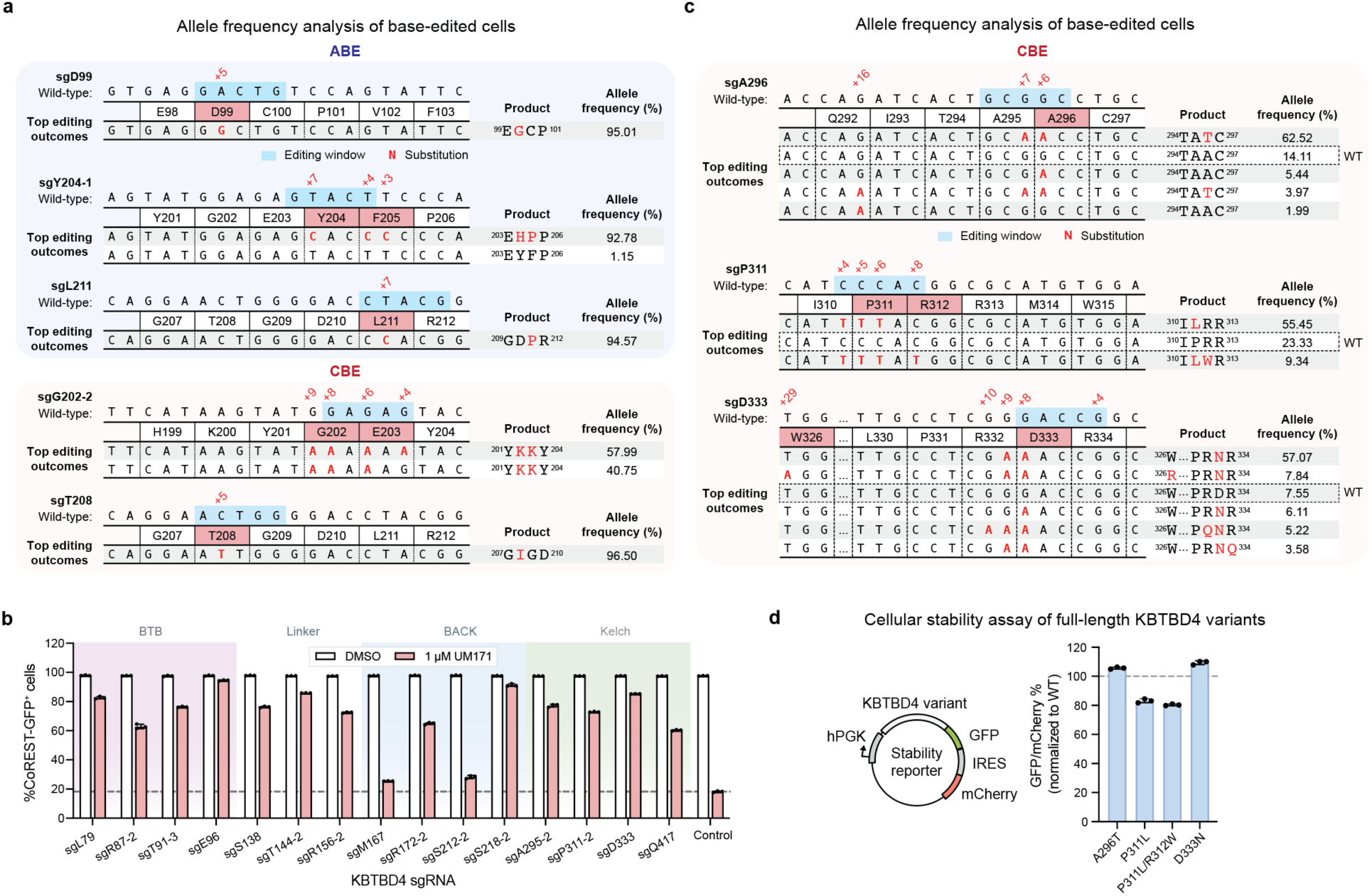
Validation of Base Edits by Individual sgRNA Transductions. **a)** Allele frequency table for 293T clonal cell lines after base editing of *HDAC1* with the indicated sgRNAs and SpG CBE. Only alleles with ≥1% allele frequency in at least one sample are shown. Protein product sequences are shown with nonsynonymous mutations in red. Genotyping was performed once (*n = 1*). **b)** Flow cytometry quantification showing GFP signal in CoREST-GFP K562 cells transduced with the indicated KBTBD4 sgRNAs, following treatment with DMSO or UM171 (1 µM) for 24 h. Bars represent mean ± s.d. of *n = 3* replicates. **c)** Base editing efficiency for selected KBTBD4 sgRNAs in K562 cells. The wild-type allele is boxed, and only alleles with ≥1% allele frequency in at least one sample are shown. Protein product sequences are shown with nonsynonymous mutations in red. Genotyping was performed once (*n = 1*). **d)** GFP to mCherry fluorescence ratio measured by flow cytometry quantification of MOLM-13 cells expressing the indicated KBTBD4-GFP reporter. Bars represent mean ± s.d. of *n = 3* replicates. KBTBD4 stability calculated as GFP/mCherry and measurements are normalized to wild-type KBTBD4 analyzed in parallel. Results in b and d are representative of two independent experiments.

**Extended Data Table 1.**
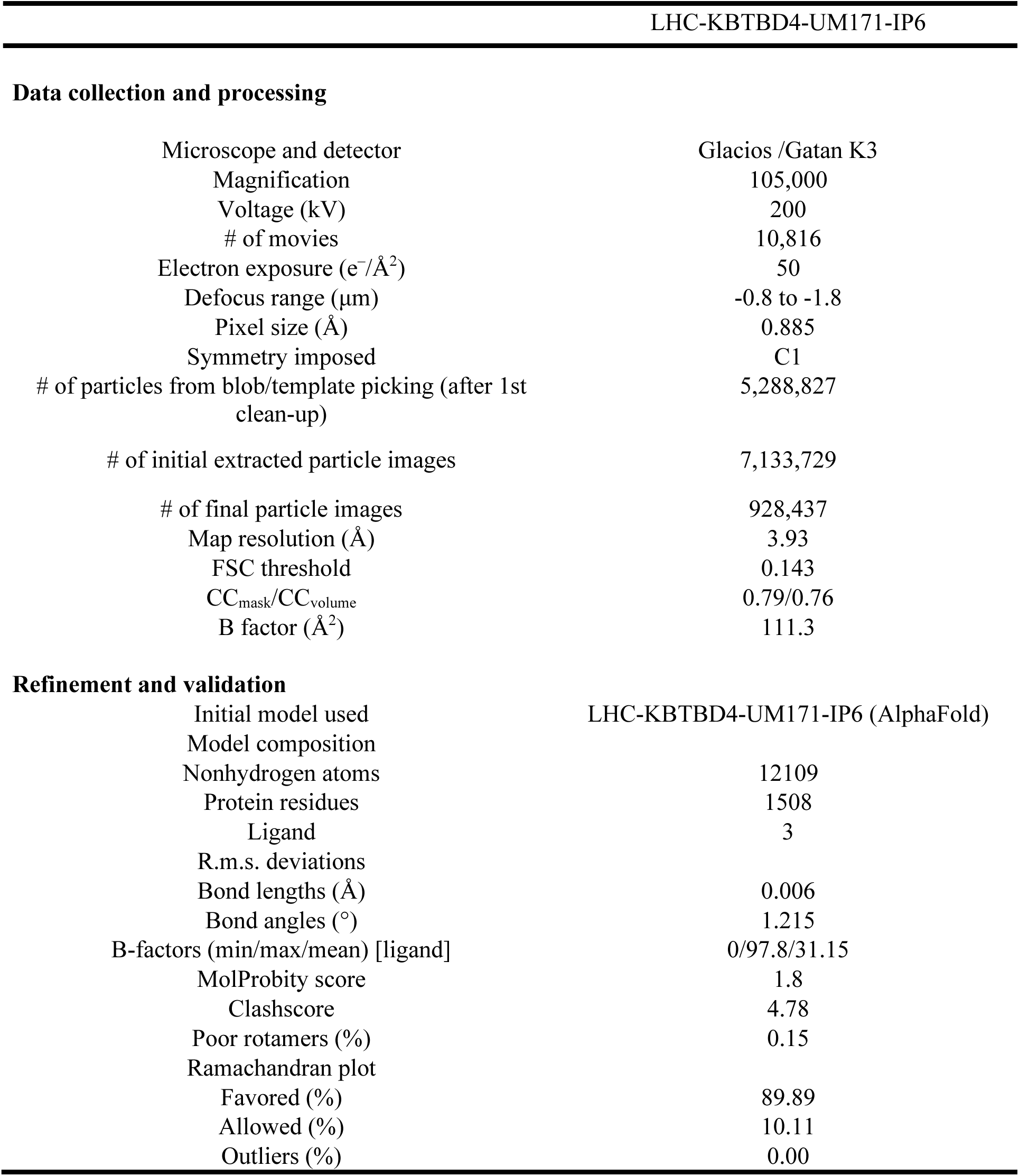
Cryo-EM data collection, refinement, and validation statistics.

## Materials and Methods

### Cell culture

MOLM-13 and SET-2 cells were a gift from M.D. Shair (Harvard University). HEK293T cells were a gift from B.E. Bernstein (Massachusetts General Hospital). MV4;11 and K562 cells were obtained from ATCC. HEK293F cells were obtained from Thermo Fisher. All mammalian cell lines were cultured in a humidified 5% CO_2_ incubator at 37 °C and routinely tested for mycoplasma (Sigma-Aldrich). RPMI1640 and DMEM media were supplemented with 100 U ml^−1^ penicillin and 100 µg ml^−1^ streptomycin (Gibco) and FBS (Peak Serum). MOLM-13, MV4;11, and K562 cells were cultured in RPMI1640 (Gibco) supplemented with 10% FBS. SET-2 cells were cultured in RPMI1640 (Gibco) supplemented with 20% FBS. HEK293T cells were cultured in DMEM (Gibco) supplemented with 10% FBS. HEK293F cells were cultured in Freestyle™ 293 Expression Medium (Thermo Fisher) shaking at 125 rpm. *Spodoptera frugiperda* (Sf9) insect cells (Expression Systems, 94-001F) were cultured in ESF921 media (Expression Systems) in a non-humidified and non-CO_2_ incubator at 27 °C shaking at 140 rpm. High Five cells were purchased from Thermo Fisher (B85502), with Grace insect medium (Thermo Fisher, 11595030) supplemented with 10% FBS (Cytiva) and 1% Penicillin-Streptomycin (Gibco), cultured at 26 °C.

### Lentiviral production

For lentivirus production, transfer plasmids were co-transfected with GAG/POL and VSVG plasmids into 293T cells using Lipofectamine™ 3000 (Thermo Fisher Scientific) according to the manufacturer’s protocol. Media was exchanged after 6 h and the viral supernatant was collected 52 h after transfection and sterile-filtered (0.45 µm). MOLM-13 and K562 cells were transduced by spinfection at 1,800 × *g* for 1.5 h at 37 °C with 5 µg mL^-1^ and 8 µg mL^-1^ polybrene (Santa Cruz Biotechnology), respectively. Where necessary, 48 h post-transduction, cells were selected with 1 µg mL^-1^ and 2 µg mL^-1^ puromycin (Thermo Fisher Scientific), respectively.

### Plasmid construction

sgRNAs were ordered as synthetic oligonucleotides (Azenta/Genewiz), annealed, and ligated into the appropriate vector: lentiCRISPR.v2 (Cas9 knockout), a gift from F. Zhang (Addgene no. 52961); pRDA_478 (Addgene no. 179096), which expresses BE3.9 (SpG), or pRDA_479 (Addgene no. 179099) which expresses ABE8e (SpG) for base editing. For individual sgRNA validation of the KBTBD4 CBE screen, sgRNAs were cloned into a pRDA_256 (Addgene no. 158581) vector containing SpG Cas9 NG PAM. Other plasmids were cloned by Gibson Assembly using NEBuilder HiFi (New England Biolabs). Cloning strains used were NEB Stable (lentiviral) and NEB 5-alpha (other plasmids) (New England Biolabs). For base editor cloning, bacterial cultures were grown at 30 °C. Final constructs were validated by Sanger sequencing (Azenta/Genewiz).

All KBTBD4 expression plasmids encoded isoform 1 (human, residues 1-518) but longer isoform 2 (residues 1-534) numbering was used. CoREST expression plasmids encoded isoform 1 (human) in either full length (residues 1-482) or various truncations. ORFs of human KBTBD4 and CoREST (mammalian expression) were obtained from Horizon Discovery. Full length MIER1 isoform 1 (human, residues 1-512) ORF was obtained from GeneCopoeia and full length RCOR2 isoform 1 (human, residues 1-523) was a gift from M.L. Suvà (Massachusetts General Hospital).

For fluorescent and stability reporter constructs, CoREST, MIER1, RCOR2, and KBTBD4 were cloned into Cilantro 2, a gift from B. Ebert (Addgene no. 74450). For transfection constructs, CoREST-FLAG and HA-KBTBD4 constructs were cloned into pcDNA3. For KBTBD4 overexpression constructs, KBTBD4 coding sequences were cloned into pSMAL mCherry, which was generated from pSMAL through introduction of an mCherry ORF into pSMAL (a gift from J. E. Dick, University of Toronto), or pFUGW-IRES-puro, which was generated from pFUGW (Addgene no. 14883) by replacing the UbC promoter-EGFP cassette with an EFS-NS-IRES-puromycin cassette. For bacmid expression, KBTBD4 was cloned into pFastbac, a gift from T. Cech. The KBTBD4 construct for structure determination was made by cloning human KBTBD4 cDNA isoform 1 into a pFastBac vector with a tandem 10X His-tag and MBP tag at the N-terminus followed by a TEV protease cutting site.

### CRISPR/Cas9-mediated genome editing

mEGFP followed by a ‘GGGSGGGS’ linker was knocked into the C-terminus of CoREST (i.e., *RCOR1*) in K562 cells. sgRNA (sgRNA: TTCAAAGCCACCAGTTTCTC) targeting the C-terminus of CoREST was cloned into a Cas9 plasmid, PX459^51^, and electroporated according to the manufacturer’s protocol (Neon™ Transfection System, Thermo Fisher Scientific) with a repair vector containing the mEGFP CDS and linker flanked by 750 bp of genomic homology sequences to either side of the CoREST C-terminus. Briefly, 2 x 10^5^ cells were washed twice with PBS and resuspended in buffer R. PX459 (0.5 µg) and the repair vector (0.5 µg) were added to the cell suspension, and electroporated at 1350 V with 10 ms pulse width for 4 pulses using the Neon^TM^ Transfection System 10 µL kit. After electroporation, cells were immediately transferred to prewarmed media. To generate single-cell clones, cells were gated to sort for the top 0.2% GFP^+^ and single-cell sorted on a MoFlo Astrios EQ Cell Sorter (Beckman Coulter), expanded, and validated by western blot and Sanger sequencing.

### Generation of knockout K562s

Lentiviral vectors carrying sgRNA (HDAC1, HDAC2) were generated by cloning appropriate sequences (HDAC1: GCACCGGGCAACGTTACGAA; HDAC2: TACAACAGATCGTGTAATGA) into pLentiCRISPR.v2 lentiviral vector. Control vector contained sgRNA targeting luciferase (sgLucA). Lentivirus was produced and K562 CoREST-GFP cells were transduced and puromycin selected as described above.

HDAC2-null and KBTBD4-null CoREST-GFP K562 clones were generated by using the Alt-R™ CRISPR-Cas9 System (IDT) to deliver ribonucleoprotein complexes containing HDAC2 KO guides (KBTBD4: GATATCTGTGAGTAAGCGGT; HDAC2: TACAACAGATCGTGTAATGA) using the Neon™ Transfection System (Thermo Fisher Scientific) according to the manufacturer’s protocol. Transfected cells recovered for 72 h before sorting for single cell clones on a MoFlo Astrios EQ Cell Sorter (Beckman Coulter). Single cell clones were validated by genotyping and immunoblotting. sgRNA and primer sequences for validation are provided in Supplementary Data 1.

### Proteomics sample preparation

MV4;11 and SET-2 (50 million cells per replicate) were treated with 1 µM UM171 or DMSO for 6 h. Cells were washed twice with ice-cold PBS, and snap-frozen in liquid nitrogen for storage at – 80 °C until use (n=3, biological replicates). Frozen cell pellets were lysed in DPBS (Fisher Scientific) supplemented with Benzonase (Santacruz Biotechnology) and Protease Inhibitor Cocktail (Roche) using a chilled bath sonicator at 4 *°*C (Q700, QSonica). Lysates were clarified by centrifugation at 300 × *g* for 3 min. Proteins were quantified by BCA assay (Thermo Fisher Scientific) and normalized to 200 µg/150 µL. 200 µg of protein was reduced with 5 mM Tris(2-carboxyethyl) phosphine hydrochloride (TCEP) (Sigma Aldrich) for 2 min and alkylated with 20 mM chloroacetamide (CAA) for 30 min at room temperature. 1000 µg of magnetic SP3 beads (1:1 hydrophobic:hydrophilic) (Cytiva) was added to each sample along with 100% LC/MS-grade ethanol (Sigma Aldrich) to reach the final concentration of 50% ethanol. Samples were then incubated for 30 min with KingFisher Flex system (Thermo Fisher Scientific) at room temperature. Beads were washed three times with 80% HPLC-grade ethanol (Sigma Aldrich) and resuspended with 150 µL of Trypsin/Lys-C (4 µg, Thermo Fisher Scientific) in 200 mM EPPS (pH 8.4)/5 mM CaCl_2_ (Sigma Aldrich), and proteins were digested overnight for 16 h at 37 *°*C. Digested peptides were dried by a Speedvac, reconstituted with 5% acetonitrile (Sigma Aldrich)/0.1% formic acid (Thermo Fisher Scientific) and desalted using EmporeTM C18 Extraction Disks (3M). Peptides were eluted with 80% acetonitrile/0.1% formic acid, dried by a Speedvac. Peptides reconstituted with 5% acetonitrile/0.1% formic acid were quantified using Quantitative Colorimetric Peptide Assay (Thermo Fisher Scientific) and 10 µg of peptides for each sample were labeled with 50 µg of TMTpro16-plex reagents (Thermo Fisher Scientific) per channel. TMT labeling was performed for 75 min with rotation at room temperature, and reaction was quenched by adding 5% hydroxylamine (Acros Organics) for 15 min, followed by addition of 10% formic acid. Samples were then pooled and dried by a Speedvac.

### High-pH reversed phase peptide fractionation

Peptides were reconstituted with 300 µl of 5% acetonitrile/0.1% formic acid. Fractionation was performed using Pierce High pH Reversed-Phase Peptide Fractionation Kit (Thermo Fisher Scientific) according to the manufacturer’s instruction. Briefly, peptide samples were fractionated with 21 increments (7.5-55% with every 2.5% increase, and 75%) of acetonitrile with 10 mM NH_4_HCO_3_. Three eluents from every 7^th^ fraction were pooled to get total seven fractions and dried by a Speedvac.

### Mass spectrometry data acquisition

Fractionated samples were reconstituted with 2% acetonitrile/0.1% formic acid and analyzed on an EASY-nLC 1200 system (Thermo Fisher Scientific) coupled to an Orbitrap EclipseTM TribridTM Mass Spectrometer (Thermo Fisher Scientific) with FAIMSpro system equipped with real-time search function. Peptides were loaded onto a trap column (Pepmap 100 C18, 3 μm particle size, 100 Å pore size, 75 μm i.d. x 150 mm length) and separated over a 140 min gradient of 5-35% acetonitrile in 0.1% formic acid and a flow rate of 300 nL/min with an analytical column (EASY-Spray C18 HPLC, 2 μm particle size, 75 µm i.d. x 500 mm length). Peptides were acquired by data-dependent acquisition (DDA) and quantified using synchronous precursor selection MS3 (DDA-SPS-MS3); briefly, peptides were ionized at 2300 V, separated by FAIMSpro (1.5 s per cycle), and scanned for MS1 analysis (resolution of 120,000; scan range of 400-1400 m/z; maximum ion injection time (IIT) 50 ms; automatic gain control (AGC) setting of 10,000). MS2 analysis was collected from collision-induced dissociation (CID, collision energy of 36%), and MS3 spectra were analyzed in the orbitrap (resolution 50,000; mass range 100-500 Da).

### Mass spectrometry data analysis

Data processing was performed in ProteomeDiscoverer (PD) v2.5 (Thermo Fisher Scientific) using the SequestHT algorithm. All raw files were submitted to search against the UniProtKB human universal database (UniProt UP000005640, downloaded May 2020) combined with the common Repository of Adventitious Proteins (cRAP, classes 1, 2, 3, and 5) and the following parameters^52^; precursor tolerance of 10 ppm, fragment ion tolerance of 0.6 Da, minimum peptide length of 6, and trypsin full digestion with zero miscleavages. Cysteine carbamidomethylation (+57.021 Da) and methionine oxidation (+15.995 Da) were set as a variable modification while lysine- and N-terminus-TMTpro modification (+304.207 Da) were set as static modifications. Peptide-spectrum matches (PSMs) were filtered to a 1% false discovery rate (FDR) using the Percolator algorithm and further for protein assignment. Reporter ion quantifier node was set with the co-isolation threshold of 50, signal-to-noise (S/N) threshold of 10, and SPS mass matches threshold of 50. Peptide abundance was normalized to total peptides. Protein ratio was calculated using PD2.5 pairwise ratio-based algorithm and Students’ t-test was used for statistical significance. Data are provided in Supplementary Data 2-3.

### Co-immunoprecipitation/mass spectrometry (Co-IP/MS)

#### Co-immunoprecipitation (Co-IP)

2 x 10^7^ SET-2 cells were washed with cold PBS and lysed on ice in lysis buffer (50 mM Tris-HCl pH 7.5, 150 mM NaCl, 1% IGPAL-CA-630) supplemented with protease and phosphatase inhibitors (Roche) and the lysates were cleared. The protein concentration was quantified as above and diluted to 1 mg/mL in lysis buffer. Antibodies were crosslinked to Protein G Dynabeads (ThermoFisher Scientific) using BS3. Crosslinked Dynabeads were washed thrice with lysis buffer and incubated with the precleared protein lysate. Immunoprecipitation was carried out overnight at 4 °C under constant rotation. Beads were washed once with wash buffer A (50 mM Tris-HCl pH 7.5, 150 mM NaCl, 0.05% IGPAL-CA-630) and twice with wash buffer B (50 mM Tris-HCl pH 7.5, 150 mM NaCl) and eluted in SDS-PAGE loading buffer and carried forward to immunoblotting as described above or mass spectrometry as described below.

#### Mass spectrometry

Beads were washed 50 mM Tris HCl (500 µL, pH 7.5, 4X) and transferred to fresh 1.7 mL Eppendorf tubes (Axygen, MCT175LC) after the second and fourth washes. Proteins were digested off beads by adding 1 mM DTT (150 µL), 5 µg/mL Trypsin (Promega), 2 M urea, 50 mM Tris HCl digestion buffer and incubating for 1 hour at 25 °C, with shaking. The supernatant was removed and saved and an additional 150 µL of digestion buffer was added and incubated for 30 min at 25 °C, with shaking. The supernatants were combined, and the beads were washed 2M Urea, 50 mM Tris HCl (150 µL, 2X), added to the saved supernatants, and transferred to fresh tubes for in-solution digestion. Proteins were reduced with 4 mM DTT, alkylated with 10 mM IAA, and digested with 0.5 µg of Trypsin overnight at 25°C, with shaking. The samples were quenched the following day with formic acid to a final concentration of 1% FA (pH < 3). Samples were desalted with C18 stage tips (2 punches) following standard protocol. Samples were loaded onto the tips and washed with 0.1% FA (100 µL, 2X) before eluting with 50% ACN, 0.1% FA (50 µL), frozen and lyophilized. Desalted samples were resuspended in 11 µL of 100 mM HEPES, pH 8.5, and labeled with 5 µL of TMT-11 reagent (Thermo Scientific, LOT# TE270748 - TD264064) at a concentration of 20 ug/µL in 100% ACN. The labeling reaction occurred at room temperature for 1 h, with shaking. Conditions were as follows: LSD1 wild type, DMSO (126C, 127N) and rabbit IgG (130C, 131N). Samples were then diluted with 5 µL of 70 mM HEPES, 30% ACN. Samples were quenched with 5% Hydroxylamine (1 µL), combined, lyophilized, and resuspended 3% ACN, 5% FA (150 µL). TMT-labeled samples were fractionated by high-pH C18 reverse phase fractionation (3 punches). 40 µg were loaded onto the stage tip and washed with 1% FA (100 µL, 2X). Following high-pH equilibration with 3% ACN, 20 mM ammonium formate (250 µL, 4X), six fractions were collected (300 µL each) at 8%, 12%, 16%, 20%, 25% and 50%. Fractions were collected in HPLC vials, were frozen and lyophilized. Each fraction was resuspended in 3% ACN/5% FA (8µL) for MS analysis (2 µL).

Fractionated samples were analyzed on an Orbitrap Q-Exactive HF-X Plus MS (Thermo Fisher Scientific) equipped with a nanoflow ionization source and coupled to a nanoflow Proxeon EASY-nLC 1000 UHPLC system (Thermo Fisher Scientific). Acquisition occurred in positive ion mode. Samples were injected on an in-house packed column (22 cm x 75 µm diameter C18 silica picofrit capillary column) heated at 50 °C. The mobile phase flow rate was 250 nL/min of 3% ACN/1% FA (solvent A) and 90% ACN/0.1% FA (solvent B). Peptides were separated using the following LC gradient: 0-6% B in 1 min, 6-30% B in 85 min, 30-60% B in 9 min, 60-90% B in 1 min, stay at 90% B for 5 min, 90-50% B in 1 min, and stay at 50% B for 5 min. Data was acquired in centroid mode for both MS1 and MS2 scans. Samples were analyzed in data dependent analysis (DDA) mode using a Top-20 method. Ion source parameters were: spray voltage 2 kV, source temperature 250 °C. Full MS scans were acquired in the m/z range 350–1800, with an AGC target 3e6, maximum IT 10 ms and resolution 60,000 (at m/z 200). MS/MS parameters were as follows: AGC target 5e4, maximum IT 105 ms, loop count 10, isolation window 0.7 m/z, NCE 31, resolution 45,000 (at m/z 200) and fixed first mass 100 m/z; unassigned and singly charged ions were excluded from MS/MS.

Raw MS data were analyzed using Spectrum Mill Proteomics Workbench (prerelease version B.06.01.202, Agilent Technologies). A trypsin specific enzyme search was performed using a 2017 uniprot human fasta file (UniProt.human.20171228.RISnrNF.553smORFs.264contams) containing 65095 entries. Peptide and fragment tolerances were at 20 ppm, minimum matched peak intensity 40% and peptide false discovery rates (FDR) were calculated to be <1% using the target-decoy approach^53^. Fixed modifications were carbamidomethylation, TMT 10 (N-term, K) and variable modifications were Acetyl (ProN-term), Oxidized methionine (M), Pyroglutamic acid (N-termQ) and Deamidation (N). Spectra with a score <4 were filtered out. Peptides were validated using the following parameters: for charge states 2-4, a FDR of 1.2 was applied to each run and for charge state 5, a FDR of 0.6 was applied across all runs. Results were further validated at the protein level and proteins with a score of 20 or higher were accepted as valid. Reporter ion correction factors, specific to the TMT batch, were applied. A protein/peptide summary was generated using the median across all TMT channels as the denominator. Shared peptides were assigned to the protein with the highest score (SGT). Calculated ratios at the protein level were imported into Protigy for normalization and features selection (https://github.com/broadinstitute/protigy). To account for variability between samples, log ratios were normalized by centering using the sample Median and scaled using the sample Median Absolute Deviation (Median MAD). Data are provided in Supplementary Data 4.

### Fluorescent degradation reporter assay

CoREST (full length and truncated), MIER1, and RCOR2 inserts were PCR-amplified with Esp3I sites and ligated into a Cilantro 2 EGFP-IRES-mCherry reporter vector by golden-gate assembly. Point mutations were introduced into coding regions using standard PCR-based site-directed mutagenesis techniques. Deletion constructs were made by PCR amplification of the appropriate regions and cloned into the Cilantro 2 vector using Gibson cloning (New England Biolabs). Lentiviral particles carrying the respective constructs in the Cilantro 2 vector were produced and used to transduce MOLM-13 cells as described above. 48 h after transduction, cells were selected with 5 µg ml^-1^ puromycin concentration for 3-5 days. The selected cells were then treated with various concentrations of UM171 or 0.1% vehicle for 6 or 24 h. GFP and mCherry fluorescence were measured on a NovoCyte 3000RYB flow cytometer (Agilent) after drug or vehicle treatment. The geometric mean of the ratio of GFP to mCherry fluorescence was calculated for each sample using the NovoExpress software (v. 1.5.0, Agilent). The ratios for the individual drug-treated samples were normalized to the ratios of the vehicle-treated samples. All degradation assays were done in triplicate and flow cytometry gating strategy is shown in **Extended Fig. 2a**.

### Immunoblotting

Cells were lysed on ice in RIPA buffer (Boston BioProducts) with 1X Halt Protease Inhibitor Cocktail (Thermo Fisher Scientific) and 5 mM EDTA (Thermo Fisher Scientific). Lysate was clarified by centrifugation and total protein concentration was measured with the BCA Protein Assay (Thermo Fisher Scientific). Samples were electrophoresed and transferred to a 0.45 μm nitrocellulose membrane (Bio-Rad). Membranes were blocked with tris-buffered saline Tween (TBST) with 5% Blotting-Grade Blocker (Bio-rad) and incubated with primary antibody: KBTBD4 (Novus Biologicals, catalog no. NBP1-88587, 1:1,000), HDAC1 (Cell Signaling Technology, catalog no. 34589, D5C6U, 1:1,000), HDAC2 (Cell Signaling Technology, catalog no. 57156, D6S5P, 1:1,000), FLAG (Sigma-Aldrich, catalog no. F1804, M2, 1:2,000), His-tag (Cell Signaling Technology, catalog no. 2365, 1:1,000), HA-tag (Cell Signaling Technology, catalog no. 3724, C29F4, 1:1,000), GAPDH (Santa Cruz Biotechnology, catalog no. sc-47724, 0411, 1:10,000), H3K9ac (Abcam, catalog no. AB32129, 1:2,000), H3 (Abcam, catalog no. AB1791, 1:2,000). Membranes were washed 3X with TBST and incubated with secondary antibody: anti-rabbit IgG HRP conjugate (Promega, catalog no. W4011, 1:20,000), anti-mouse IgG HRP conjugate (Promega, catalog no. W4021, 1:40,000), goat anti-rabbit IgG HRP conjugate (Cell Signaling Technology, catalog no. 7074, 1:2,000). Following 3X washes with TBST, immunoblots were visualized using SuperSignal West Pico PLUS or SuperSignal West Femto chemiluminescent substrates (Thermo Fisher Scientific).

### Co-immunoprecipitation

#### In K562 cells

FLAG-KBTBD4 was cloned into pFUGW-IRES-puro and stably expressed in CoREST-GFP K562 cells via lentiviral transduction followed by puromycin selection, as described above. Cells were pre-treated with either 10 µM Cpd-60 (12 h), 10 µM SAHA (1 h), or vehicle, then treated with 1 µM MLN4924 for 3 h then 5 µM UM171 or vehicle for 1 h. Cells were washed twice with cold PBS and flash frozen. Co-IP was performed as described below.

#### In HEK293T cells

HEK293T cells were transfected with 3 µg pcDNA3 HA-KBTBD4 plasmid and 3 µg pcDNA3 CoREST-FLAG (full-length or truncated) using PEI MAX transfection reagent (Polysciences) according to the manufacturer’s protocol. 48 h after transfection, cells were treated with 1 µM MLN4924 for 3 h then 1 µM UM171 or vehicle for 1 hour. Cells were washed twice with cold PBS and flash frozen. Co-IP was performed as described below.

Cells were thawed, lysed on ice in lysis buffer (25 mM Tris-HCl pH 7.5, 150 mM NaCl, 1% NP-40 alternative) supplemented with cOmplete™, EDTA-free Protease Inhibitor Cocktail (Sigma-Aldrich), and the lysates were cleared. The protein concentration was quantified as above and diluted to 1 mg ml^−1^ in lysis buffer with 1 µM UM171 or DMSO. Supernatants were immunoprecipitated overnight at 4 °C with 25 µL Pierce anti-HA magnetic beads (Thermo Fisher Scientific). Beads were washed six times with lysis buffer, eluted in SDS-PAGE loading buffer, and carried forward to immunoblotting as described above.

### Protein expression and purifications

Recombinant human KBTBD4 for biochemical and biophysical analyses was purified from Sf9 insect cells. cDNAs for human KBTBD4 protein was cloned into the pFastBac donor vector and the recombinant baculovirus was constructed using the Bac-to-Bac protocol and reagents (Thermo Fisher Scientific). KBTBD4 construct was tagged on the N-terminus with 6×His cleavable by TEV protease. This plasmid was used to prepare baculovirus according to standard protocols (Bac-to-Bac Baculovirus Expression System, Thermo Fisher). Detection of gp64 was used to determine baculovirus titer (Expression Systems). For expression, Sf9 cells were grown to a density of 1-2 x10^6^ cells/mL and infected with KBTBD4 baculovirus. The cells were incubated for 72 h (27 °C, 120 × *g*), harvested and then frozen with liquid nitrogen for future purification. Cells were resuspended in lysis buffer (50 mM Tris-HCl, pH 8.0 cold, 500 mM NaCl, 1 mM TCEP, 10% glycerol, 15 mM imidazole) supplemented with 1% NP-40, 1 mM PMSF, and cOmplete™, EDTA-free Protease Inhibitor Cocktail (Sigma-Aldrich) and sonicated. Lysates were clarified by centrifugation at 100,000 × *g* for 30 min and incubated with His60 Ni Superflow affinity resin (Takara). Resin was washed with lysis buffer containing a stepwise gradient of 15-50 mM imidazole, followed by elution using lysis buffer with 250 mM imidazole. Eluate was exchanged into storage buffer (50 mM Tris-HCl, pH 8.0 cold, 150 mM NaCl, 1 mM TCEP, 10% glycerol) using an Econo-Pac 10DG desalting column (Bio-Rad) and further purified by size exclusion chromatography using a Superdex 200 10/300 GL column (GE Healthcare). The purity of the recombinant protein was verified by SDS-PAGE and fractions with 90-95% purity were pooled and stored at –80 °C.

Recombinant human KBTBD4 used in cryo-EM structure determination was purified from *Trichoplusia ni* High Five insect cells. cDNAs for human KBTBD4 was cloned into the pFastBac donor vector and the recombinant baculovirus was constructed using the Bac-to-Bac protocol and reagents (Thermo Fisher Scientific). KBTBD4 construct was tagged on the N-terminus with 10xHis and MBP tag cleavable by TEV protease. This plasmid was used to prepare baculovirus according to standard protocols (Bac-to-Bac Baculovirus Expression System, Thermo Fisher). For expression, the monolayer High Five cells were grown to about 80% confluency and infected with KBTBD4 baculovirus. The cells were incubated for 72 h (26 °C), harvested, and then frozen with liquid nitrogen for future purification. Cells were resuspended in lysis buffer (50 mM Tris-HCl, pH 8.0 cold, 150 mM NaCl, 1 mM TCEP, 20 mM imidazole) supplemented with 1 mM PMSF, 10 µM Leupeptin, 0.5 µM Aproptinin and 1 µM Pepstatin A and sonicated. Lysate was clarified by centrifugation at 100,000 × *g* for 30 min and incubated with amylose affinity resin (New England BioLabs). Resin was washed with lysis buffer, followed by elution using lysis buffer with 10 mM maltose. Eluate was cut with Tobacco Etch Virus protease overnight, followed by the prepacked anion exchange column (GE Healthcare) to get rid of the protease and further purified by size exclusion chromatography using a Superdex 200 10/300 GL column (GE Healthcare). The purity of the recombinant protein was verified by SDS-PAGE and fractions with 90–95% purity were pooled and stored at –80°C.

Recombinant full length HDAC1 was purified from Sf9 insect cells^54^. cDNA for human HDAC1 with C-terminal FLAG tag was cloned into the pFastBac donor vector using standard PCR-based site-directed mutagenesis techniques and the recombinant baculovirus was constructed using the Bac-to-Bac protocol and reagents (Thermo Fisher Scientific). Detection of gp64 was used to determine baculovirus titer (Expression Systems). For expression, Sf9 cells were grown to a density of 1-2 x10^6^ cells/mL and infected with HDAC1 baculovirus at a MOI of 5. The cells were incubated for 72 h (27 °C, 120 × *g*), harvested and then frozen with liquid nitrogen for future purification. Cells were resuspended in lysis buffer (20 mM Tris-HCl, pH 7.4, 10% glycerol, 0.5 M KCl, 5 mM MgCl2, 0.1% NP40, 1 mM PMSF supplemented with cOmplete™, EDTA-free Protease Inhibitor Cocktail (Sigma-Aldrich) and incubated for 45 min at 4 °C with stirring to allow for lysis. The lysate was clarified at 100,000 × *g* for 30 min and incubated with FLAG resin (Anti-Flag M2 affinity gel, Sigma) for 2 h. The resin was washed 4 times with lysis buffer and one final time with the same buffer containing 150 mM KCl. The FLAG tagged protein was eluted in lysis buffer with 150 mM KCl supplemented with 200 µg/mL FLAG peptide for 1 h at 4 °C. The purity of the complex was verified by SDS-PAGE and stored at –80 °C.

Recombinant LSD1-CoREST (LC) complex was comprised of LSD1 (aa 151-852) and CoREST (aa 305-482). LSD1 (aa 151-852) was cloned into a pET15b vector (gift from P. A. Cole) containing an N-terminal 6×His-tag using NEBuilder HiFi DNA Assembly Master Mix (NEB E2621L). The LSD1 constructs were expressed in BL21-CodonPlus (DE3)-RIPL competent *E. coli* and after plating a single colony was cultivated in 2xYT with 100 mg/L ampicillin at 37 °C and expression was induced at OD600 of 1.0 by adding 0.3 mM isopropyl β-D-thiogalactoside (IPTG) and grown for 5 h at 25 °C. CoREST(aa 305-482) was expressed from a pGEX vector (gift from A. Mattevi). The plasmid was transformed into BL21-CodonPlus (DE3)-RIPL *E.coli* cells and after plating a single colony was cultivated in LB media with 100 mg/L ampicillin at 37 °C and expression was induced at OD600 of 0.8 by adding 0.25 mM isopropyl β-D-thiogalactoside (IPTG) and grown overnight at 17 °C. The cells were pelleted by centrifugation at 4,000 × *g* for 30 min and stored at –80 °C prior to purification. All purification steps were performed at 4 °C. Pellets of CoREST and LSD1 were resuspended in lysis buffer (50 mM NaH_2_PO_4_ pH 8.0, 300 mM NaCl, 5% glycerol, 7.5 mM imidazole supplemented with PMSF, DNAse and EDTA-free Roche protease inhibitor cocktail) in a weight ratio of 1:1.5, respectively. Cells were disrupted by sonication, clarified by centrifugation and passed through nickel affinity resin as before. The eluent was then loaded onto GST resin equilibrated in GST affinity buffer (50 mM NaH_2_PO_4_ pH 8.0, 300 mM NaCl, 5% glycerol, 1 mM DTT, 1 mM EDTA) and the GST-tag was cleaved on the resin after incubation with GST-PreScission protease (APEXBIO) overnight at 4 °C. The protein was eluted by washing the column with GST affinity buffer, concentrated and subsequently gel-filtered on a Superdex 200 10/300 GL column equilibrated in storage buffer as before. The purity of the complex was verified by SDS-PAGE and fractions with 90-95% purity were pooled and stored at –80 °C.

Recombinant LSD1-CoREST-HDAC complex was comprised of full length LSD1 (UniProt ID: O60341) or LSD1 (Δ77-86), full length HDAC1 (UniProt ID: Q13547) and N-terminally truncated CoREST (aa 84-482) (UniProt ID: Q9UKL0) or N-terminal Cys CoREST^17^. The pcDNA3 vector was used to create plasmids encoding the different proteins. The CoREST constructs contained an N-terminal 10xHis-3xFLAG tag followed by a TEV protease cleavage site. The constructs for ternary complex were co-transfected into suspension-grow HEK293F cells (ThermoFisher Scientific) with polyethylenimine (PEI) (Sigma) and harvested after 48 h. Cells were resuspended in lysis buffer (50 mM HEPES, pH7.5, 100 mM KCl, 5% glycerol, 0.3% Triton X-100, 1X Roche EDTA-free Complete Protease Inhibitor cocktail) and sonicated. Lysate was clarified by centrifugation at 12,000 rpm for 30 min, and the supernatant was incubated with Anti-Flag M2 affinity gel (Sigma). The affinity gel was washed twice with lysis buffer and twice with SEC buffer (50 mM HEPES, pH 7.5, 50 mM KCl, 0.5 mM TCEP) followed by the incubation with TEV protease overnight at 4 °C. The complex was further purified by size exclusion chromatography using a Superose 6 10/300 column (GE Healthcare). The purity of the complex was verified by SDS-PAGE and fractions with 90-95% purity were pooled and supplemented with 5% glycerol and stored at –80 °C.

### Fluorescein labeling of LHC

The fluorescein labeling of the LSD1-CoREST-HDAC1 complex was purified as described above. A Cys point mutagenesis was conducted next to the TEV protease cleavage site of N-terminally truncated CoREST for the ligation reaction with NHS-fluorescein^55^. A 2 mM NHS-fluorescein was incubated with 500 mM mercaptoethanesulfonate (MESNA) in the reaction buffer (100 mM HEPES, pH 7.5, 50 mM KCl, 1 mM TCEP) for 4 h at room temperature in the dark for transesterification. The LSD1-CoREST-HDAC1 complex purified by FLAG M2 affinity gel was washed with reaction buffer and incubated with TEV protease for 5 h at 4 °C. The complex was then mixed with 500 µL of the fluorescein/MESNA solution to make a final concentration of 0.5 mM fluorescein and 125 mM MESNA. The mixture was incubated for 48 h at 4 °C in the dark. The complex was desalted by a Zeba spin desalting column (7 kDa MWCO) and further purified by size exclusion chromatography using a Superose 6 10/300 column (GE Healthcare). Fluorescein-labeling efficiency was analyzed by SDS-PAGE and fluorescence gel imaging (Amersham Typhoon FLA 9500, Cytiva). The purity of the complex was verified by SDS-PAGE and fractions with 90-95% purity were pooled and supplemented with 5% glycerol and stored at –80 °C.

### Fluorescence polarization measurements

#### Titration of KBTBD4

Recombinant WT KBTBD4 was diluted to 15 µM in a one-to-one mixture of ligand buffer (50 mM Tris-HCl, pH 8.0 cold, 150 mM NaCl, 1 mM TCEP, 10% glycerol) and LHC buffer (20 mM HEPES pH 7.5, 1 mM TCEP, 2 mg/mL BSA, 0.1% Tween-20, +/- 100 µM InsP_6_) containing 10 nM **JL1** with or without 20 nM recombinant LHC, HDAC1, or LC. This was aliquoted in triplicate into a black 384-well plate (Corning), followed by twofold serial dilution in assay buffer containing 10 nM **JL1** (final volume, 25 µL). The plate was incubated at room temperature for 1 h and read (1,700 ms integration) using a SpectraMax i3x with a rhodamine fluorescence polarization cartridge and SoftMax Pro software (Molecular Devices). Wells containing only assay buffer were used for background subtraction. The G-factor was adjusted to set the polarization of assay buffer with 10 nM **JL1** and 200 nM LHC only to a reference value of 27 mP. Curves were fitted to the sigmoidal, 4PL model in GraphPad Prism 9.

#### Titration of SAHA

Recombinant WT KBTBD4 (5 µM) and recombinant LHC (20 nM) were diluted to in a one-to-one mixture of ligand buffer (50 mM Tris-HCl, pH 8.0 cold, 150 mM NaCl, 1 mM TCEP, 10% glycerol) and LHC buffer (20 mM HEPES pH 7.5, 1 mM TCEP, 2 mg/mL BSA, 0.1% Tween-20, 100 µM InsP_6_) containing 10 nM **JL1** and 10 µM SAHA. This was aliquoted in triplicate into a black 384-well plate (Corning), followed by twofold serial dilution in assay buffer containing 10 nM **JL1**, recombinant WT KBTBD4 (5 µM), and recombinant LHC (20 nM) (final volume, 25 µL). The plate was incubated at room temperature for 1 h and read (1,700 ms integration) using a SpectraMax i3x with a rhodamine fluorescence polarization cartridge and SoftMax Pro software (Molecular Devices). Wells containing only assay buffer were used for background subtraction. The G-factor was adjusted to set the polarization of assay buffer with 10 nM **JL1** and 200 nM LHC only to a reference value of 27 mP. Curves were fit to the sigmoidal, 4PL model in GraphPad Prism 9.

### Microscale thermophoresis measurements

MST assays were performed with a Monolith NT.115 (NanoTemper) using the Nano BLUE mode. The exciting laser power was set at 50% and MST power was set to Medium. K_D_ values were calculated using MO.analysis (v2.3) software with a quadratic equation binding K_D_ model shown below.

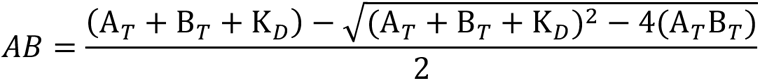

#### Titration of UM171

Fluorescein-labeled LHC (200 nM) was titrated with UM171, in the absence or presence of wild-type KBTBD4 (200 nM) in a one-to-one mixture of ligand buffer (50 mM Tris-HCl, pH 8.0 cold, 150 mM NaCl, 1 mM TCEP, 10% glycerol) and LHC buffer (20 mM HEPES pH 7.5, 1 mM TCEP, 2 mg mL^-1^ BSA, 0.1% Tween-20, 100 µM InsP_6_) at 23 °C. UM171 (up to 100 µM) was prepared with a twofold serial dilution. Then LHC with or without WT KBTBD4 was added, mixed well, and incubated for 10 min for equilibration before transferring to MST premium capillaries. Prism 9 was used to fit the data to a four-parameter dose-response curve.

#### Titration of KBTBD4

Fluorescein-labeled LHC (200 nM) was titrated with WT KBTBD4, in the absence or presence of DMSO or UM171 (50 µM) in the MST binding assays at 23 °C. WT KBTBD4 (up to 11.7 μM) was prepared with a twofold serial dilution for titrating with fluorescein-LHC. 50 μM UM171 or equivalent amount of DMSO were added. Then LHC (or LC) at a final concentration of 200 nM was added, mixed well, and incubated for 10 min for equilibration before transferring to MST premium capillaries. Prism 9 was used to fit the data to a four-parameter dose-response curve.

### Histone H3K9ac synthesis

The depsipeptide as Fmoc-Thr(O*t*Bu)-glycolic acid was synthesized based on a reported two-step protocol^56^. Then, H3K9ac(aa1-34) with a sequence as ARTKQTARKS-TGGKAPRKQL-ATKAARKSAP-A-**TOG**-G was synthesized via standard solid phase peptide synthesis (SPPS) and purified by reversed phase HPLC. The Fmoc-protected amino acids were purchased from Novabiochem except for Fmoc-Lys(Ac)-OH (EMD Millipore 852042). F40 sortase was expressed and purified as reported previously, and bacterial expression and purification of *X. laevis* globular H3 (gH3; amino acids 34-135 C110A) were performed also following a previous protocol^56^. Then, the F40 sortase catalyzed histone H3 ligation reaction was carried out between the H3K9ac (aa1-34, please note the C-terminal residue is extruded) peptide and the gH3. The reaction mixture was purified by ion-exchange chromatography to obtain pure semisynthetic histone H3K9ac (C110A) characterized by MALDI-TOF MS as reported previously^57^.

### Octamer refolding and nucleosome reconstitution

146 bp Widom 601 DNA was prepared by previously reported methods used for the nucleosome reassembly^58^. Bacterial expression and purification of *X. laevis* core histones H2A, H2B and H4 were then carried out, followed by assembly of the histone octamer and refolding as previously reported^59^. The octamer was purified by size exclusion chromatography using a Superdex 200 10/300 GL column (GE Healthcare) and was used for nucleosome assembly with 146 bp 601 Widom DNA as reported previously^60^. The final mixture was subjected to HPLC purification (Waters, 1525 binary pump, 2489 UV-Vis detector) with a TEKgel DEAE ion exchange column to purify the final nucleosome product. The purified nucleosome containing H3K9ac was analyzed by native TBE-gel with EtBr staining, as well as SDS-PAGE gel and then Western blot analysis by anti-H3K9ac antibody^60^.

### Analysis of LHC complex deacetylation of acetylated nucleosome

The general deacetylation assay was set up as reported previously^61^. The LHC complex was diluted into the pH 7.5 reaction buffer containing 50 mM HEPES, 100 mM KCl, 0.2 mg/mL BSA, and 100 μM InsP_6_ to a final concentration of 90 nM. After the addition of KBTBD4 to a final concentration of 300 nM and/or UM171 (in final 10% DMSO) to a final concentration of 10 μM, the solution was pre-incubated for 15 min at ambient temperature. After chilling on ice for 3 min, the deacetylation reaction was initiated with the addition of H3K9ac nucleosome to a final concentration of 100 nM, and all the reaction solutions were incubated for 120 min at 37 °C. Different aliquots were taken at time points of 0, 30 min, 60 min, 90 min, and 120 min. Each aliquot was quenched with an SDS-loading buffer containing 20 mM EDTA, and was heated at 95 °C for 3 min. After running SDS-PAGE and iBlot transfer to nitrocellulose membranes, western blot analysis was performed with anti-H3K9ac primary antibody (Abcam, AB32129, 1:2000 dilution), followed by the Goat anti-Rabbit secondary antibody (Cell Signaling Technology, 7074S, 1:2000 dilution). Western blot analysis with anti-H3 (Abcam, AB1791, 1:2000 dilution) was used as the loading control. Imaging analysis with chemiluminescence on GeneSys was quantified using ImageJ software^57^. All intensity values were fit to a single-phase exponential decay curve with constrain Y0=1, Plateau=0 (GraphPad Prism Ten). Each plotted point represents 2 replicates for the kinetic parameter V/[E] calculation.

### HDAC1/2 activity assays

Recombinant HDAC1 (BPS Bioscience 50051) or HDAC2 (BPS Bioscience 50002) were diluted to 6 nM (1.2×) into buffer containing 50 mM HEPES, pH 7.5, 100 mM KCl, 0.5 mg/mL BSA, 0.001% Tween-20 and 25 μL added to wells of a white, 384-well microtiter plate (Corning 3572). Test compounds were added in serial dilution (1:2 titration, 15-point, c_max_ = 10 μM) using a D300 digital dispenser (Hewlett-Packard), and allowed to equilibrate for 1 h at room temperature. Then, 5 μL of 6× MAZ1600 HDAC substrate^27^ was added (final HDAC1/2 concentration 5 nM; final MAZ1600 concentration 18 μM) and deacetylase activity was allowed to proceed for 45 min at room temperature. Next, 5 μL of 7× developer solution was added (150 nM trypsin + 40 μM LBH589 final concentrations) and the plate was incubated for 30 min at room temperature. 7-Amino-4-methyl coumarin fluorescence was measured on a Tecan Spark plate reader: 350/20 nm excitation, 460/10 nm emission. The assay floor (background) was defined with the 10 μM LBH589 dose, and the assay ceiling (top) was defined via a no-inhibitor control. Data were background-corrected, normalized, and Prism 9 was used to fit the data to a four-parameter dose-response curve.

### TR-FRET measurements

Unless otherwise noted, experiments were performed in white, 384-well microtiter plates (Corning 3572) in 30 μL assay volume, or white, 384-well low-volume microtiter plates (PerkinElmer 6008280). TR-FRET measurements were acquired on a Tecan SPARK plate reader with SPARKCONTROL software version V2.1 (Tecan Group Ltd.), with the following settings: 340/50 nm excitation, 490/10 nm (Tb), and 520/10 nm (FITC, AF488) emission, 100 μs delay, 400 μs integration. The 490/10 nm and 520/10 nm emission channels were acquired with a 50% mirror and a dichroic 510 mirror, respectively, using independently optimized detector gain settings unless specified otherwise. The TR-FRET ratio was taken as the 520/490 nm intensity ratio on a per-well basis.

### Ternary complex measurements by TR-FRET

#### Titration of UM171

Recombinant WT 6×His-KBTBD4 (40 nM), fluorescein-labeled LSD1-CoREST-HDAC complex (40 nM), and CoraFluor-1-labeled anti-6×His IgG (20 nM)^31^ were diluted into a one-to-one mixture of ligand buffer (50 mM Tris-HCl, pH 8.0, 150 mM NaCl, 1 mM TCEP, 10% glycerol) and LHC buffer (20 mM HEPES, pH 7.5, 1 mM TCEP, 2 mg/mL BSA, 0.1% Tween-20, 100 μM InsP_6_), with or without 100 μM SAHA, and 10 μL added to wells of a white, 384-well low volume microtiter plate (PerkinElmer 6008280). UM171 was added in serial dilution (1:3 titration, 10-point, c_max_ = 10 μM) using a D300 digital dispenser (Hewlett-Packard), and allowed to equilibrate for 1 h at room temperature before TR-FRET measurements were taken. Data were background-corrected from wells containing no UM171. Prism 9 was used to fit the data to a four-parameter dose-response curve.

#### Titration of UM171 and InsP_6_

Recombinant WT 6×His-KBTBD4 (40 nM), fluorescein-labeled LSD1-CoREST-HDAC complex (40 nM), and CoraFluor-1-labeled anti-6×His IgG (20 nM)^31^ were diluted into were diluted into a one-to-one mixture of ligand buffer (50 mM Tris-HCl, pH 8.0, 150 mM NaCl, 1 mM TCEP, 10% glycerol) and LHC buffer (20 mM HEPES, pH 7.5, 1 mM TCEP, 2 mg/mL BSA, 0.1% Tween-20) and 10 μL added to wells of a white, 384-well low volume microtiter plate (PerkinElmer 6008280). UM171 was added in serial dilution (1:10 titration, 5-point, c_max_ = 10 μM) and InsP_6_ was added in serial dilution (1:10 titration, 6-point, c_max_ = 100 μM) using a D300 digital dispenser (Hewlett-Packard), and allowed to equilibrate for 1 h at room temperature before TR-FRET measurements were taken. Data were background-corrected from wells containing no UM171 and no InsP_6_. Prism 9 was used to fit the data to a four-parameter dose-response curve.

#### Titration of Fluorescein-labeled LSD1-CoREST-HDAC complex

Recombinant WT 6×His-KBTBD4 (20 nM, 2×) and CoraFluor-1-labeled anti-6×His IgG (10 nM, 2×)^31^ were diluted into LHC buffer, with or without 10 μM UM171, and 5 μL added to wells of a white, 384-well low volume microtiter plate (PerkinElmer 6008280). Serial dilutions of fluorescein-labeled LSD1-CoREST-HDAC complex (1:2 titration, 10-point, c_max_ = 1,000 nM, 2×) were prepared in ligand buffer and 5 μL added to wells of the same plate (final volume 10 μL, final 6×His-KBTBD4 concentration 10 nM, final CoraFluor-1-labeled anti-6×His IgG concentration 5 nM, fluorescein-labeled LSD1-CoREST-HDAC complex c_max_ 500 nM). The plate was allowed to equilibrate for 1 h at room temperature before TR-FRET measurements were taken. Data were background-corrected from wells containing no 6×His-KBTBD4. Prism 9 was used to fit the data to a four-parameter dose-response curve.

### *In vitro* ubiquitination assay

The ubiquitination assays were set up similarly as previously reported^62^. Reactions were performed at 37 °C in a total volume of 20 µL. The reaction mixtures contained 5 mM ATP, 100 μM WT ubiquitin, 100 nM E1 protein, 2 μM E2 protein, 0.5 μM neddylated RBX1-CUL3, 0.5 µM WT KBTBD4 (unless otherwise indicated), 10 µM UM171/DMSO with 25 mM Tris-HCl (pH 7.5), 20 mM NaCl, 10 µM InsP_6_, and 2.5 mM MgCl2 as reaction buffer. Substrate fluorescein-LHC at 0.5 µM was preincubated with everything except E1 in the reaction mixture at 37 °C for 5 min prior to adding E1 to initiate the reaction. Reactions were quenched at the indicated time points by adding SDS loading buffer containing reducing agent β-mercaptoethanol. The reaction samples were resolved on SDS-PAGE gels and analyzed by Colloidal Blue staining and western blots.

### Base editor scan

The sgRNA libraries were designed as described previously to include all sgRNAs (NG protospacer-adjacent motif) targeting exonic and flanking ±30 bp into the intronic regions of canonical isoforms of KBTBD4 (ENST00000430070.7) and HDAC1 (ENST00000373548.8), excluding those with TTTT sequences. Negative (nontargeting, intergenic) and positive (essential splice site) controls were included^63^. The library was synthesized as an oligonucleotide pool (Twist Biosciences) and cloned into pRDA_478 and pRDA_479 following published workflows. Lentivirus was produced and titered by measuring cell counts after transduction and puromycin selection. Cells were transduced with library lentivirus at a multiplicity of infection <0.3 and selected with puromycin for 5 days. Cells were then expanded and split into three replicate subcultures and treated with DMSO or 1 µM UM171. After 24 h, cells were sorted on a MoFlo Astrios EQ Cell Sorter (Beckman Coulter), collecting top 10% GFP^+^ and unsorted (GPF^±^) cells. Genomic DNA was isolated using the QIAamp DNA Blood Mini kit, and sgRNA sequences were amplified using barcoded primers, purified by gel extraction, and sequenced on an Illumina MiSeq as previously described^21,64^. At all steps, sufficient coverage of the library was maintained in accordance with published recommendations.

Data analysis was performed using Python (v.3.9.12) with Biopython (v.1.78), Pandas (v.1.5.1), SciPy package (v.1.10.0) and NumPy (v.1.23.4). sgRNA enrichment was calculated as previously described^21,64^. Briefly, sequencing reads matching each sgRNA were quantified as reads per million, increased by a pseudocount of 1, log2-transformed, normalized to the plasmid library and replicate-averaged. Sorted GFP^+^ abundances were normalized to unsorted abundances. The mean value for non-targeting controls was subtracted to calculate the final enrichment value for each sgRNA; this value is referred to as the normalized log_2_(fold-change sgRNA enrichment). sgRNAs that zero counts in the plasmid libraries were excluded from further analysis.

sgRNAs with scores >4 s.d. above or below the mean of intergenic negative controls were considered ‘enriched’ or ‘depleted,’ respectively. sgRNAs targeting KBTBD4 and HDAC1 were classified based on expected editing outcome, assuming any C or A within the editing window (protospacer +4 to +8) of cytidine and adenosine base editors, respectively, is converted to T. sgRNAs were placed in one of six mutually exclusive classes: in order of assignment priority, (1) nonsense; (2) missense; (3) silent; (4) UTR-intronic; (5) non-editing (no Cs and/or As); (6) negative controls (does not target gene). Library sgRNA annotations and base editor scanning data are provided in Supplementary Data 5-12. Scatter and line plots were generated using matplotlib (v3.7.1).

### Linear clustering analysis

Per-residue sgRNA enrichment scores were estimated as previously described^43^. Briefly, LOESS regression was performed on using the ‘lowess’ function of the statsmodels package (v.0.13.5) in Python (v.3.9.12) with a 20 AA sliding window (‘frac = (20 AA/L)’, where L is the total length of the protein), and ‘it = 0’ to fit observed log_2_(fold-change sgRNA enrichment), hereafter sgRNA enrichment score, as a function of amino acid position. Only sgRNAs that are predicted to result in missense mutations were used. For amino acid positions that were not targeted by sgRNAs, enrichment scores were interpolated by performing quadratic spline interpolation on the LOESS output scores using the ‘interp1d’ function of the SciPy package (v.1.10.0).

To assess statistical significance of the resulting clusters, we simulated a null model of random sgRNA enrichment scores. Amino acid positions of sgRNAs were kept fixed while sgRNA enrichment scores were randomly shuffled, and per-residue enrichment scores were recalculated by performing LOESS regression and interpolation on the randomized sgRNA enrichment scores for each of 10,000 permutations. Empirical p values were calculated for each amino acid by comparing its observed resistance score to the null distribution of random resistance scores. Empirical p values were adjusted using the Benjamini–Hochberg procedure to control the false discovery rate to ≤0.05. Finally, linear clusters were called by identifying all contiguous intervals of amino acids with adjusted p values P ≤0.05. For plotting, adjusted p values were increased by a pseudocount of 10-4, log_10_-transformed and multiplied by -1.

### Genotyping

Genomic DNA was purified using the QIAamp DNA Blood Mini (Qiagen). We subjected 100 ng of DNA to a first round of PCR (25-28 cycles, Q5 hot start high-fidelity DNA polymerase (New England Biolabs)) to amplify the locus of interest and attach common overhangs. 1 µL of each PCR product was amplified in a second round of PCR (8 cycles) to attach barcoded adapters. Primer sequences are provided in Supplementary Table. Final amplicons were purified by gel extraction (Zymo) and sequenced on an Illumina MiSeq. Data were processed using CRISPResso2^65^ using the following parameters: --quantification_window_size 20 -- quantification_window_center -10 --plot_window_size 20 --exclude_bp_from_left 0 -- exclude_bp_from_right 0 --min_average_read_quality 30 --n_processes 12 --base_editor_output.

### Generation of HDAC1 mutant clones

sgRNAs enriched in the base editing screens were ordered as synthetic oligonucleotides (Azenta/Genewiz), annealed, and ligated into either pRDA_478 or pRDA_479. The plasmids were transfected into HEK293T cells using Lipofectamine™ 3000 (Thermo Fisher Scientific) according to the manufacturer’s protocol. 48 h post-transduction, cells were selected with 2 µg mL^-1^ puromycin (Thermo Fisher Scientific) for 3 days, then sorted for single cell clones on a BD FACSAria Cell Sorter (BD Biosciences). Single cell clones were validated by genotyping and stability of mutants was assessed by immunoblotting. sgRNA sequences and annotations, and primer sequences used for genotyping are provided in Supplementary Data 13-14.

### Single guide validation in K562

sgRNAs enriched in the KBTBD4 CBE screen were ordered as synthetic oligonucleotides (Azenta/Genewiz), annealed, and ligated into SpG Cas9 NG PAM into the pRDA_256 plasmid. Lentivirus was produced as described above and transduced into CoREST-GFP K562. After puromycin selection, cells were harvested and validated by genotyping. sgRNA sequences and annotations are provided in Supplementary Data 13.

### Degradation assay of KBTBD4 mutants

K562 KBTBD4-null CoREST-GFP cells were generated as described above. KBTBD4 overexpression constructs were cloned into pSMAL mCherry and point mutations were introduced into coding regions using standard PCR-based site-directed mutagenesis techniques. Lentiviral particles carrying the overexpression constructs were produced and used to transduce K562 KBTBD4-null CoREST-GFP cells as described above. 48 h after transduction, cells were treated with 1 µM UM171, 1 µM MLN4924, 10 µM HDACi (SAHA, CI-994, or RBC1HI), or 0.1% vehicle for 24 h. GFP^+^% was measured for mCherry^+^ cells in each condition (**Extended Data Fig. 2c**).

### Cryo-EM sample preparation and data collection

To assemble the complex of KBTBD4-UM171-LHC for cryo-EM study, the individually isolated KBTBD4 protein and co-expressed LHC complex were mixed in stoichiometric amounts with 1 μM UM171 added and subsequently applied to the Superose6 increase gel filtration column (Cytiva) in a buffer containing 40 mM HEPES, pH 7.5, 50 mM KCl, 100 µM InsP_6_ and 0.5 mM TCEP (tris(2-carboxyethyl)phosphine). The isolated complex was then crosslinked with 37.5 mM glutaraldehyde at room temperature for 6 min and quenched the reaction with 1 M Tris-HCl pH 8.0. The crosslinked sample was snap-frozen for future use.

To prepare grids for cryo-EM data collection, a QuantiFoil Au R0.6/1 grid (Electron Microscopy Sciences) was glow discharged for 30 sec at 20 mA with a glow discharge cleaning system (PELCO easiGlow). 3.0 μL of the purified KBTBD4-UM171-LHC complex at 0.7 mg mL^-1^ was applied to a freshly glow-discharged grid. After incubating in the chamber at 10 °C and 100% relative humidity, grids were blotted for 3 sec with a blotting force of zero, then immediately plunge-frozen in liquid ethane using a Vitrobot Mark IV system (Thermo Fisher Scientific). Data collection was carried out on a FEI Titan Glacios transmission electron microscope (Thermo Fisher Scientific) operated at 200 kV at the Arnold and Mabel Beckman Cryo-EM Center of the University of Washington. Automation scheme was implemented using the SerialEM^66^ software using beam-image shift^67^ at a nominal magnification of 105 K, resulting a physical pixel size of 0.885 Å. The images were acquired on a K3 camera direct detector. The dose rate was set to 10 e^-^/Å^2^ S, and the total dose of 50 electrons per Å^2^ for each image were fractionated into 99 EER (Electron-event representation) frames. Data were collected in four sessions with a defocus range of 0.8-1.8 μm. In total, 6,839 movies were collected.

### Image processing and 3D reconstruction

In total, 10,816 movies were collected and imported into CryoSPARC^68^ followed by patch motion correction and patch CTF estimation. 10,637 micrographs were kept after filtering the micrographs with CTF parameters and manual inspection. Blob picker job in CryoSPARC was able to pick 7,133,729 particles, which were further extracted and subjected to 2D classification. After five rounds of cleaning by 2D classification, 928,437 particles were selected and subjected to ab-initio reconstruction. Subsequently, all the particles were used for heterogenous refinement. After one extra round of cleaning up by heterogenous refinement, 186,315 particles from good reconstruction were selected to get re-extracted without Fourier cropping. the homogenous refinement and non-uniform refinement^69^, help reach an overall resolution of 3.38 Å. To better optimize map for the KELCH-repeat domain, a soft mask focused on KELCH domain was applied to local refinement, ending up with a further improved resolution to 3.93 Å. More details about the data processing can be found in **Extended Data Fig. 4**.

### Model building and refinement

The initial structural models of the KBTBD4 dimer and the HDAC1-CoREST-ELM-SANT1 complex was predicted with AlphaFold-Multimer in Google ColabFold^70^. Because the overall model did not fit into the 3.93 Å cryo-EM map very well, the structural models of KBTBD4 BTB-BACK domain, KELCH-repeat domain, and HDAC1-CoREST were separately fit into the cryo-EM map using UCSF Chimera^71^. The resulting model was subsequently rebuilt in Coot^72^ based on the protein sequences and the EM density and was further improved by real-space refinement in PHENIX^73,74^. The structure figures were made using PyMOL^75^.

## Data availability

Coordinates and density map of the KBTBD4-UM171-LHC-InsP_6_ complex are deposited with the PDB and the Electron Microscopy Data Bank (EMDB) under the accession code 8VOJ and EMD-43386. All other data are available in the manuscript and the source data files.

