## Supplementary Note for "Asymmetric Engagement of Dimeric CRL3^KBTBD4^ by the Molecular Glue UM171 Licenses Degradation of HDAC1/2 Complexes"

**Supplementary Note 1 | Synthetic procedure**

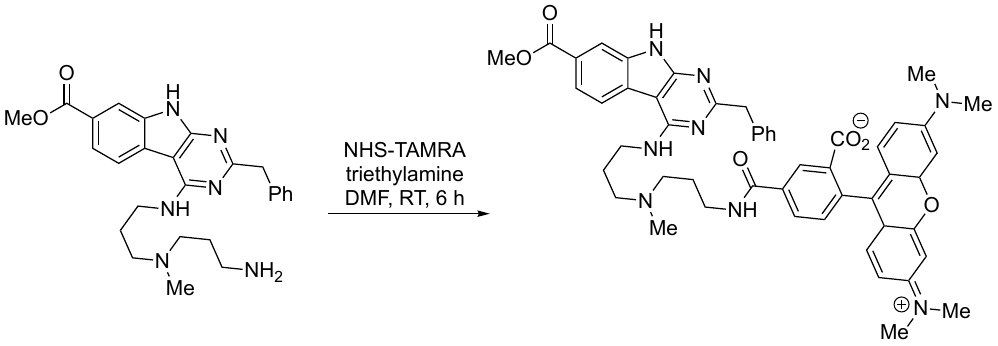

***JL1***

Methyl 4-((3-((3-aminopropyl)(methyl)amino)propyl) amino)-2-benzyl-9H-pyrimido-4,5-bindole-7-carboxylate (0.0120 g, 0.0261 mmol) was dissolved in DMF (0.5 mL) and triethylamine (10.8 uL, 0.0783 mmol), then TAMRA-NHS (0.0137 g, 0.0261 mmol) was added. The reaction mixture stirred for 6 h at RT. After evaporation to dryness, the residue underwent purification via C18 silica gel column chromatography, yielding U84-TAMRA (60%, 0.0137 g) as a purple solid. ^1^H NMR (400 MHz, METHANOL-*D*_4_) δ 8.59 (s, 1H), 8.49 (bs, 1H), 7.95 (s, 2H), 7.89 (d, *J* = 8.2 Hz, 1H), 7.47 (d, *J* = 8.2 Hz, 1H), 7.38 (d, *J* = 7.7 Hz, 2H), 7.29 - 7.25 (m, 4H), 7.20 - 7.12 (m, 2H), 7.05 (d, *J* = 9.3 Hz, 2H), 6.70 (d, *J* = 7.5 Hz, 2H), 6.43 (s, 2H), 4.09 (s, 2H), 3.79 (s, 3H), 3.52 – 3.47 (m, 4H), 3.21 - 2.96 (m, 18H), 2.70 (s, 3H), 2.03 (dm, *J* = 42.1 Hz, 4H). ^13^C NMR (101 MHz, METHANOL-D4) δ 169.60, 168.75, 168.11, 161.93, 158.68, 158.43, 158.38, 158.15, 140.46, 137.21, 136.77, 136.53, 133.14, 133.04, 132.28, 131.35, 130.40, 130.05, 129.93, 129.76, 129.58, 129.50, 129.40, 127.46, 126.59, 125.05, 122.84, 121.10, 114.91, 114.59, 113.33, 97.10, 95.52, 55.25, 55.05, 52.54, 46.72, 40.74, 39.71, 38.52, 38.05, 25.91, 25.66. HRMS-ESI [M-H]: 871.3940 and observed: 871.3962.

**Supplementary Note 2 | NMR Spectra**

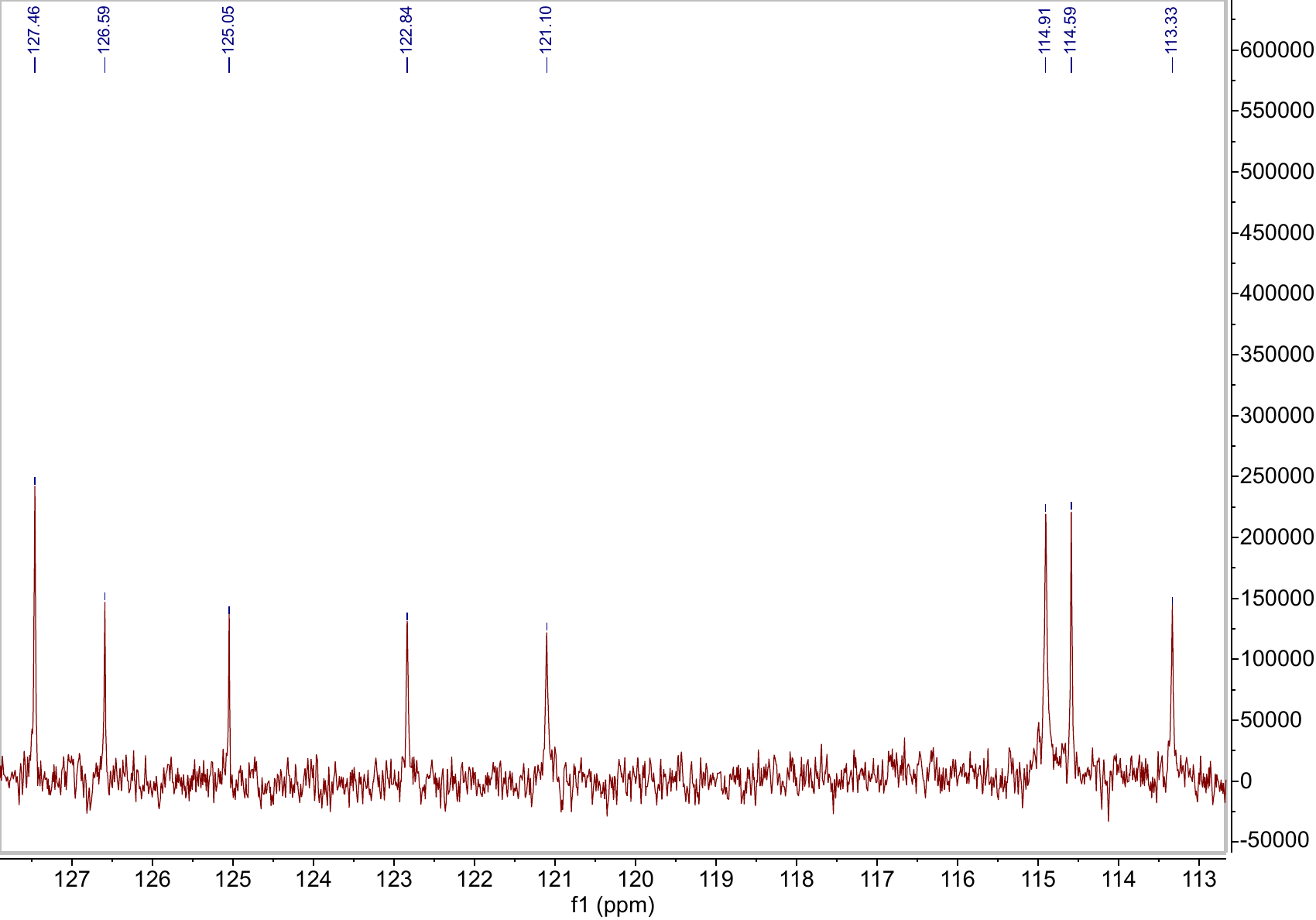

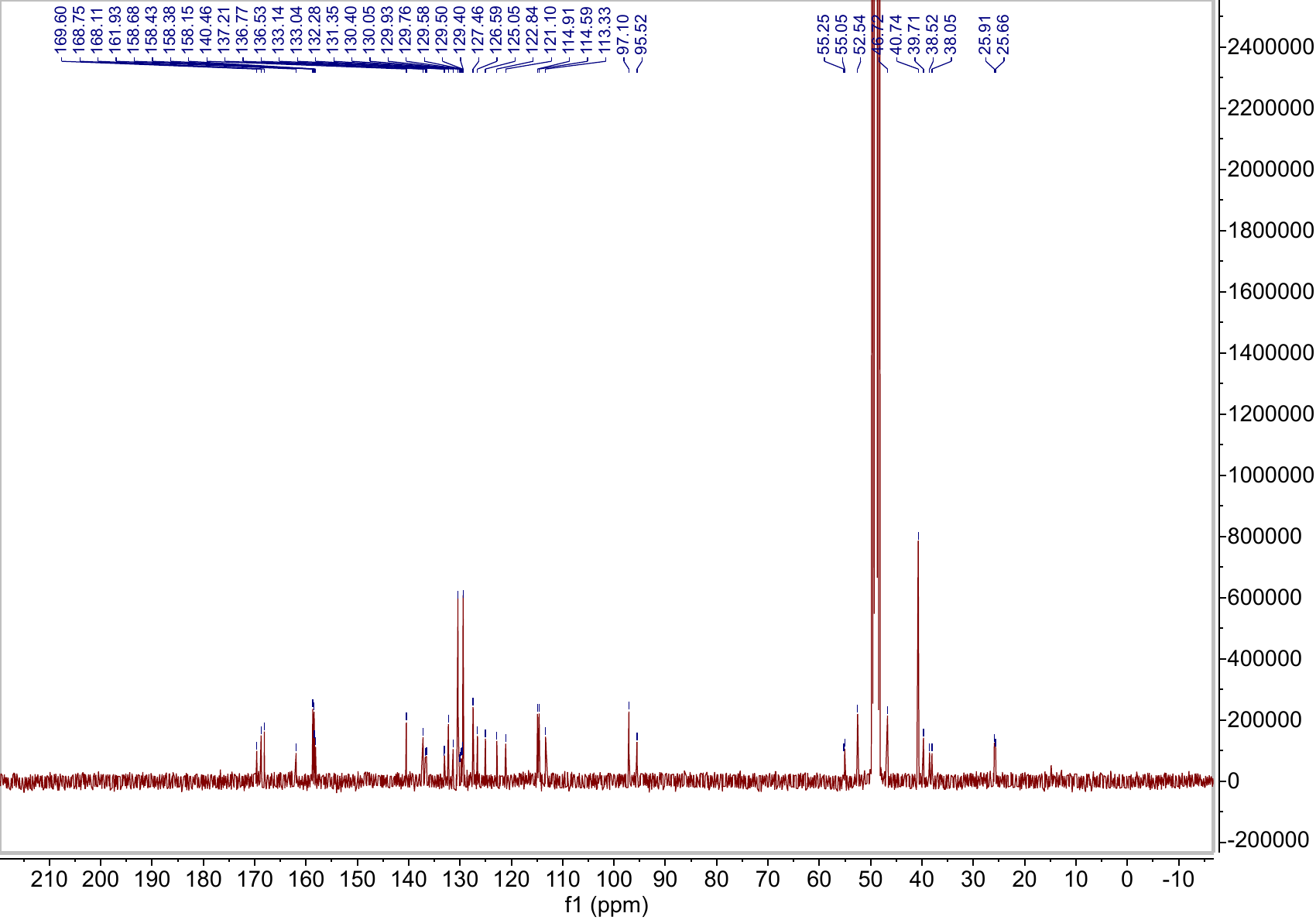

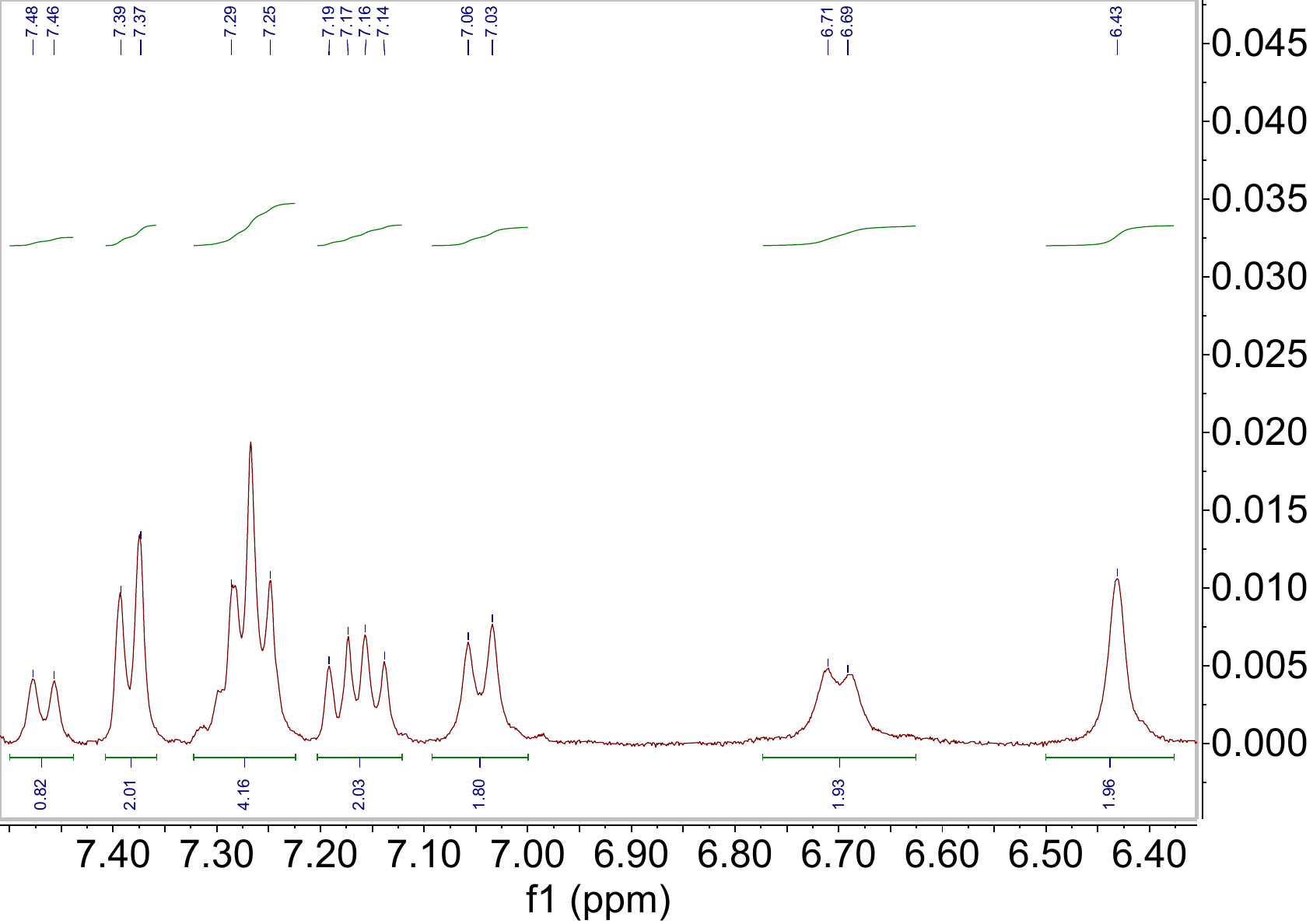

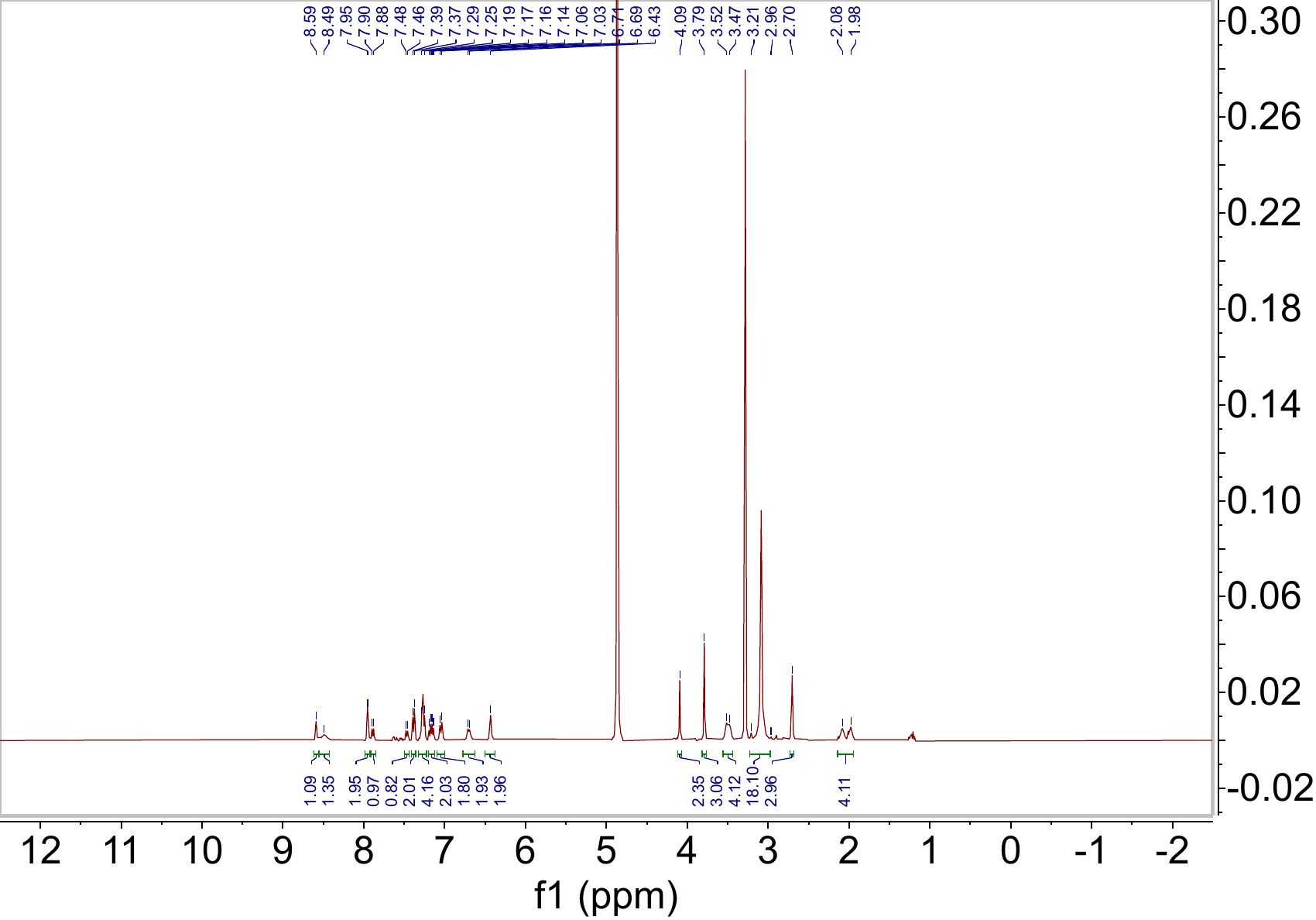
